# ATR protects centromere identity by promoting DAXX association with PML nuclear bodies

**DOI:** 10.1101/2022.09.26.509554

**Authors:** Isabelle Trier, Yoon Ki Joo, Elizabeth Black, Lilian Kabeche

## Abstract

Centromere protein A (CENP-A) defines centromere identity and nucleates kinetochore formation for mitotic chromosome segregation. Here, we show that *Ataxia telangiectasia and Rad3-related* (ATR) kinase, a master regulator of the DNA damage response, protects CENP-A occupancy at interphase centromeres in a DNA damage-independent manner. As ATR localizes to promyelocytic leukemia nuclear bodies (PML NBs) in unperturbed cells, we hypothesized that ATR protects CENP-A occupancy by regulating the localization of the histone H3.3 chaperone and PML NB component, DAXX. Indeed, we found that ATR inhibition reduces DAXX association with PML NBs, resulting in the DAXX-dependent loss of CENP-A from interphase centromeres. Lastly, we demonstrate that CENP-A occupancy is not restored until G1 of the following cell cycle, leading to increased mitotic chromosome segregation defects. These findings demonstrate a novel mechanism by which ATR protects centromere identity and genome stability.

## Introduction

Centromeres are epigenetically defined by the presence of nucleosomes containing the histone H3 variant, centromere protein A (CENP-A) (McKinley and Cheeseman, 2016; Musacchio and Desai, 2017). CENP-A nucleosomes provide a base for the hierarchical formation of the kinetochore, which links chromosomes to spindle microtubules for mitotic chromosome segregation (Cheeseman and Desai, 2008). Thus, CENP-A nucleosomes must be maintained throughout the cell cycle to build functional mitotic kinetochores. Consequently, CENP-A loss from centromeres leads to increased chromosome mis-segregation events and aneuploidy (Régnier et al., 2005), a deviation from the diploid karyotype associated with genome instability and tumorigenesis.

*Ataxia Telangiectasia and Rad3-related* (ATR) kinase is a master regulator of the DNA damage response (DDR) with diverse functions in protecting genome stability (Yazinski and Zou, 2016). ATR coordinates the cellular response to single-stranded DNA breaks and replication stress (Buisson et al., 2015), but DNA damage-independent roles have also emerged for this pleiotropic kinase. In mitosis, ATR localizes to centromeres, where it promotes proper kinetochore-microtubule attachments in a DNA damage-independent manner (Kabeche et al., 2018). During interphase and in the absence of DNA damage, ATR is enriched at promyelocytic leukemia nuclear bodies (PML NBs) (Barr et al., 2003), which house a diverse array of regulatory proteins. However, ATR’s role at PML NBs in unperturbed cells is unknown.

While many proteins transiently associate with PML NBs in response to specific stimuli, proteins involved in genome maintenance and chromatin remodeling are regularly associated with PML NBs (Corpet et al., 2020). Accordingly, PML NBs have been termed “nuclear depots” that coordinate the sequestration and release of these proteins to regulate chromatin maintenance (Morozov et al., 2012). The histone chaperones Death Domain Associated Protein 6 (DAXX) and alpha-thalassemia/mental retardation X-linked (ATRX) are considered to be “constitutive” components of these bodies (Ishov et al., 1999; Tang et al., 2004). Notably, DAXX and ATRX also localize to interphase centromeres, where they mediate the replication-independent deposition of H3.3-containing nucleosomes to regulate centromere transcription and heterochromatin formation (Morozov et al., 2012; Morozov et al., 2017). In contexts of CENP-A overexpression, DAXX is implicated in the ectopic deposition of CENP-A at chromosome arms (Athwal et al., 2015; Nye et al., 2018; Shrestha et al., 2021). Interestingly, DAXX is a putative target of ATR phosphorylation in response to genotoxic stress (Stokes et al., 2007). Thus, we hypothesized that ATR also regulates DAXX activity in the absence of DNA damage to modulate its association with PML NBs. This suggests that ATR may protect genome stability by regulating the sequestration of PML NB components in the absence of DNA damage.

Here, we show that basal ATR kinase activity protects CENP-A occupancy at interphase centromeres in a DNA damage-independent manner. CENP-A loss after acute ATR inhibition is DAXX-dependent, and ATR activity promotes DAXX association with PML NBs. Finally, we demonstrate that CENP-A occupancy at centromeres remains depleted until G1 of the following cell cycle, despite the recovery of ATR activity following inhibitor wash-out. This sustained loss of CENP-A leads to increased mitotic chromosome segregation errors. Together, these data demonstrate a novel, DNA damage-independent role for ATR in protecting centromere function and genome stability.

## Results and Discussion

### ATR kinase activity protects CENP-A occupancy at interphase centromeres

Previous work has shown that DNA damage-independent ATR kinase activity during mitosis promotes proper chromosomes segregation through the destabilization of incorrect kinetochore-microtubule attachments (Kabeche et al., 2018). Further, basal ATR activity primes the cellular response to replicative stress through the surveillance of replication forks (Yin et al., 2021) This demonstrates that ATR may possess other non-canonical functions in protecting genome stability. However, little else is known regarding DNA damage-independent ATR activity during interphase. We therefore asked whether basal ATR kinase activity may regulate the integrity of interphase centromeres, which are epigenetically defined by the presence of CENP-A-containing nucleosomes (McKinley and Cheeseman, 2016). To test this, we treated human osteosarcoma cells (U2OS) with ATR inhibitors for 1 hour (ATRi) and measured changes to CENP-A protein levels. Treatment of U2OS cells with two structurally distinct ATR inhibitors, VE-821 and AZ20, resulted in a robust decrease in a direct target of ATR kinase activity, phosphorylated serine 33 of replication protein A subunit 2 (pRPA2 S33) (Olson et al., 2006), but did not affect total cellular or nuclear CENP-A protein levels as measured by whole-cell western blotting and immunofluorescence microscopy for CENP-A (Fig. S1A-B). However, ATR inhibition decreased maximum nuclear CENP-A intensity by approximately 20% (Fig. 1A-B), suggesting that ATR kinase activity does not regulate total CENP-A levels, but may promote proper CENP-A occupancy at interphase centromeres.

**Figure 1:**
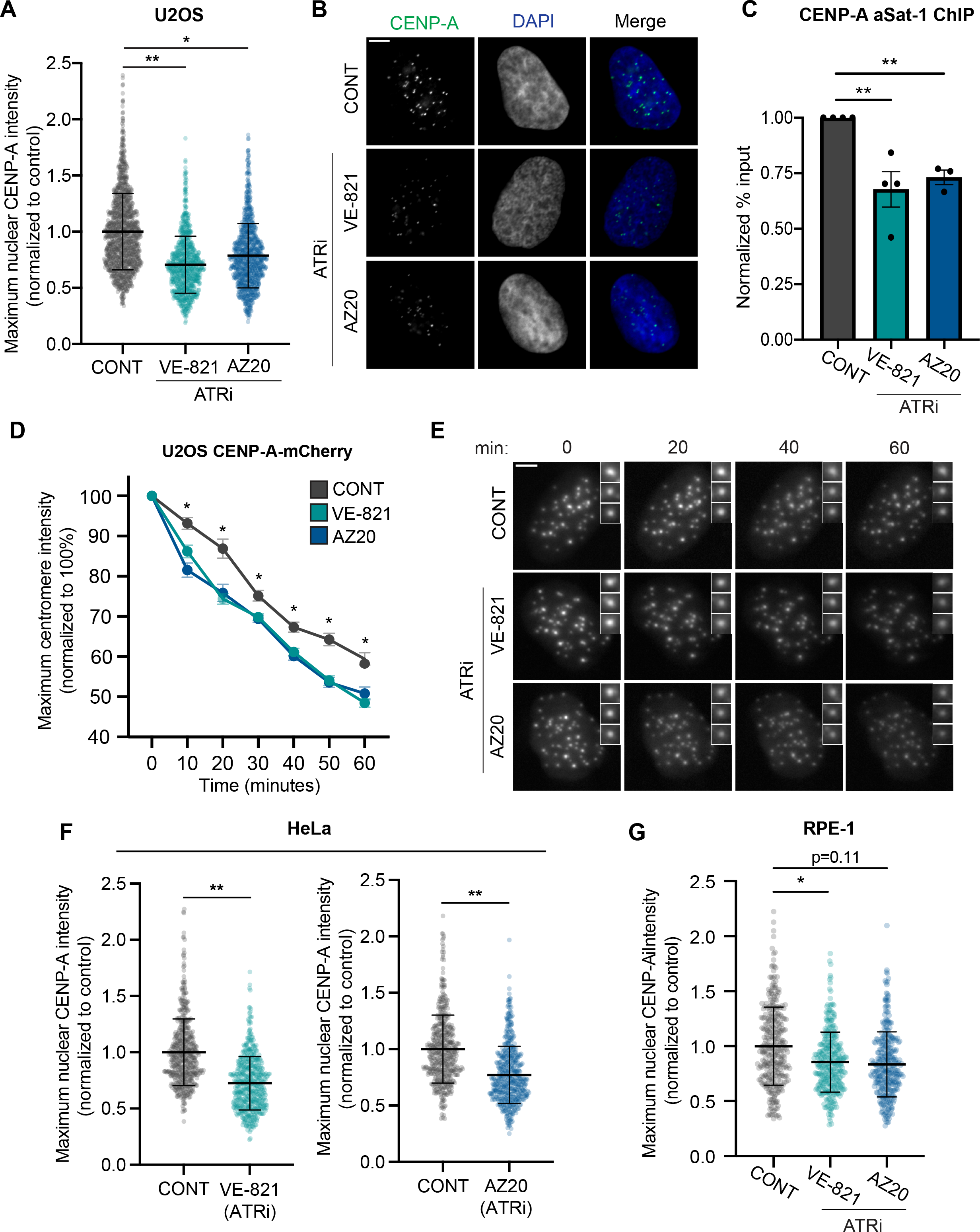
ATR kinase activity protects CENP-A occupancy at interphase centromeres. **(A-B)** *ATR inhibition reduces maximum nuclear CENP-A intensity*. (A) Quantification of CENP-A intensity in Asynchronous U2OS cells that were untreated (CONT) or treated for 1 hour with 10μM VE-821 or 10μM AZ20 (ATRi). (>180 cells per condition; n=3 biological replicates). Error bars represent mean ±SD. (B) Representative images of cells stained for CENP-A (green) and DNA (DAPI; blue). **(C)** *ATR inhibition reduces CENP-A occupancy at interphase centromeres*. Quantitative PCR of CENP-A chromatin immunoprecipitation in asynchronous U2OS cells treated as in (A). Centromeres were amplified using primers for the core α-satellite sequences of chromosomes 1, 5 and 19 (αSat-1). Error bars represent mean ±SEM. **(D-E)** *CENP-A loss with ATR inhibition is rapid*. (D) Asynchronous U2OS cells stably expressing CENP-A-mCherry were treated as in (A). The intensity of the 3 brightest CENP-A foci were quantified and averaged per cell per time point and normalized to 100% at time = 0 minutes. (10 cells per condition, n = 1 biological replicate). Error bars represent mean ±SEM. (E) Representative images of U2OS CENP-A-mCherry cells quantified in (D). Inset depicts the 3 brightest foci quantified for that cell. **(F-G)** *ATR inhibition induces CENP-A loss in HeLa and RPE-1 cells*. Quantification of maximum nuclear CENP-A intensity in asynchronous HeLa (F) and RPE-1 (G) cells treated as in (A). (>130 HeLa cells per condition and > 100 RPE-1 cells per condition; n = 3 biological replicates). Error bars represent mean ±SD. *p ≤ 0.05, **p ≤ 0.01, two-tailed t-test of replicate averages. Scale bars for all panels = 5 μm.

To verify this, we performed chromatin immunoprecipitation (ChIP) using asynchronous U2OS cells to measure CENP-A occupancy at centromere-specific alpha-satellite DNA. Using two distinct primer pairs (See STAR methods), we observed a similar decrease in CENP-A occupancy in cells treated with both VE-821 and AZ20 for 1 hour (Fig. 1C; Fig. S1C). In contrast, we did not detect a significant change in histone H2A occupancy at centromeres (Fig. S1D), suggesting that ATR inhibition leads to the targeted loss of CENP-A-containing nucleosomes without affecting total nucleosome occupancy at centromeres. Moreover, this data validates maximum nuclear CENP-A intensity as an appropriate measurement to capture changes in CENP-A occupancy at interphase centromeres.

Next, we sought to characterize the temporal dynamics of CENP-A loss from centromeres with ATR inhibition. To accomplish this, we stably infected U2OS cells with CENP-A-mCherry for live-cell imaging. Following treatment with both VE-821 and AZ20, we measured the maximum intensity of the three brightest CENP-A-mCherry foci in each cell over a 1-hour period. Although photo-bleaching led to a steady decrease in CENP-A-mCherry intensity for all conditions, cells treated with ATR inhibitors exhibited a statistically significant decrease in intensity within 10 minutes compared to control cells (Fig. 1D-E). This is consistent with our observation that ATR inhibitors act very rapidly, reducing phosphorylation of ATR’s downstream effector kinase in the DNA damage response, Chk1, in as little as 10 minutes (Fig. S1E). This demonstrates that ATR inhibition induces the rapid loss of CENP-A nucleosomes from interphase centromeres.

To determine whether this effect is seen in multiple cellular contexts, we also measured maximum CENP-A intensity using immunofluorescence microscopy in the cervical cancer cell line, HeLa, and the non-cancerous retinal pigment epithelial cell line, RPE-1. Both cell lines were sensitive to ATR inhibition with both VE-821 and AZ20, resulting in a similar loss of CENP-A from interphase centromeres (Fig. 1F-G). This demonstrates that CENP-A loss with ATR inhibition is a phenomenon observed in both cancerous and non-cancerous cell lines, suggesting a conserved role for ATR kinase in protecting centromere identity.

### CENP-A loss from centromeres with acute ATR inhibition is cell cycle phase- and DNA damage-independent

CENP-A nucleosome occupancy is tightly regulated in a cell cycle-dependent manner to ensure that centromere position and function is preserved across generations of cells. CENP-A deposition in G1 is uncoupled from its dilution across sister chromatids in S phase and from increased CENP-A transcription and translation in G2 (Jansen et al., 2007; Shelby et al., 2000; Silva et al., 2012). Thus, the decrease in centromeric CENP-A occupancy following ATR inhibition may result from phase-specific perturbations to CENP-A deposition, dilution, or expression. To address this, U2OS cells treated with ATR inhibitors were co-treated with EdU, a thymidine analog incorporated into replicating DNA, allowing for the identification of G1, S, and G2 cells according to DAPI and EdU fluorescence signal (Fig. S2A). Treatment of U2OS cells with VE-821 led to a small, albeit significant increase in the proportion of G2 cells compared to control cells. However, the proportion of cells in G1 and S phase were not significantly altered, and treatment with AZ20 had no effect on cell cycle distribution (Fig. 2A). Notably, ATR inhibition with either VE-821 or AZ20 led to a consistent decrease in maximum nuclear CENP-A intensity throughout interphase (Fig. 2B-C). This suggests that the mechanism by which ATR protects CENP-A occupancy at centromeres is independent of cell cycle phase. Taken together with the live-cell imaging experiments of U2OS CENP-A-mCherry cells (Fig. 1D-E), these results suggest that ATR inhibition leads to rapid CENP-A eviction from centromeres throughout interphase.

**Figure 2:**
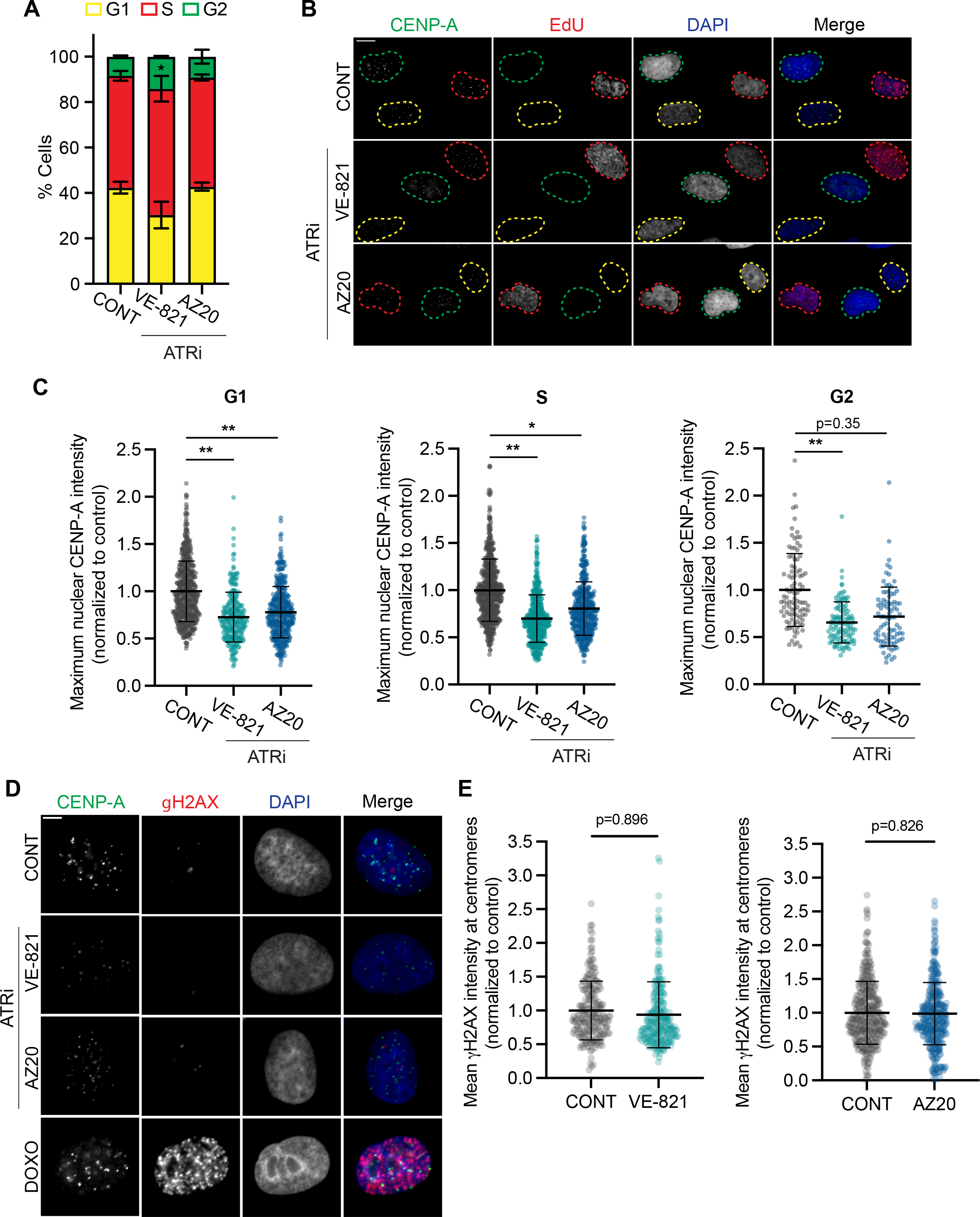
CENP-A loss from centromeres with acute ATR inhibition is cell cycle phase- and DNA damage-independent. **(A)** *Acute ATR inhibition does not impact cell cycle distribution*. Asynch. U2OS cells were untreated (CONT) or treated for 1 hour with 10μM VE-821 or 10μM AZ20 (ATRi). All cells were treated for 30 min with 10 μM EdU (>150 cells per condition; n = 3 biological replicates). Error bars represent mean ±SEM. **(B-C)** *ATR inhibition reduces nuclear CENP-A intensity in G1, S, and G2*. (B) Representative images of asynchronous U2OS cells treated as in (A). Cells were stained for CENP-A (green), EdU (red), and DNA (DAPI; blue). Nuclei are outlined according to cell cycle phase. Scale bar = 10 μm. (C) Quantification of maximum nuclear CENP-A intensity of U2OS cells sorted into their distinct cell cycle phases. (>50 G1 and S phase cells per condition, >25 G2 phase cells per condition; n = 3 biological replicates). Error bars represent mean ±SD. **(D-E)** *Acute ATR inhibition does not increase DNA damage*. (B) Representative images of asynchronous U2OS cells that were untreated (CONT) or treated for 1 hour with 10μM VE-821, 10μM AZ20, or 0.5 mM Doxorubicin (DOXO). Cells were stained for CENP-A (green), γH2AX (red), and DNA (DAPI; blue). Scale bar = 5 μm. (E) Quantification of mean γH2AX intensity at centromere foci marked by anti-centromere antibody in asynchronous (asynch.) U2OS cells that were treated as in (A). (>50 cells per condition; n = 3 biological replicates). Outliers were removed according to the ROUT (Q=1%) method. Error bars represent mean ±SD. *p ≤ 0.05, **p ≤ 0.01, two-tailed t-test of replicate averages.

DNA breaks induced by laser irradiation have been shown to recruit CENP-A nucleosomes (Zeitlin et al., 2009), and CENP-A mis-localization has also been observed in response to genotoxic stress (Hédouin et al., 2017). Due to the prominent role of ATR in the DNA damage response pathway, we assessed whether acute ATR inhibition leads to increased DNA damage. U2OS cells were treated for 1 hour with VE-821, AZ20, or Doxorubicin (DOXO), a DNA intercalator and inhibitor of topoisomerase II. As expected, treatment with Doxorubicin increased nuclear staining of phosphorylated H2AX (γH2AX; Fig. 2D), a marker of DNA damage (Mah et al., 2010), and increased total cellular γH2AX as measured by whole-cell western blotting (Fig. S2B). Alternatively, acute treatment with ATR inhibitors did not increase nuclear γH2AX signal (Fig. 2D; Fig. S2B-C), nor did ATR inhibition increase γH2AX staining at centromeres (Fig. 2E). These data are consistent with previous work demonstrating that short-term ATR inhibition does not induce significant DNA damage (Buisson et al., 2015; Kabeche et al., 2018). Further, this finding demonstrates that eviction of CENP-A nucleosomes from centromeres is unlikely to be caused by increased DNA damage at centromeres or mis-localization to sites of DNA damage in the context of acute ATR inhibition.

While acute ATR inhibition did not increase γH2AX signal in cells, we nonetheless sought to determine if CENP-A occupancy at centromeres is protected through activation of the DNA damage response pathway. In response to single-stranded DNA lesions, ATR phosphorylates its downstream effector kinase, Chk1, to induce cell cycle arrest and promote DNA repair (Liu et al., 2000). In parallel, Ataxia Telangiectasia Mutated (ATM) coordinates the cellular response to double-stranded DNA breaks (Marechal and Zou, 2013). As such, a decrease in CENP-A occupancy at centromeres with Chk1 and ATM inhibition would indicate a broad role for the DNA damage response pathway in regulating centromere identity. To test this, we treated U2OS cells with the Chk1 inhibitor, SCH 90076 (Chk1i) or the ATM inhibitor KU-55933 (ATMi) for 1 hour and measured maximum nuclear CENP-A intensity. Surprisingly, neither Chk1 nor ATM inhibition led to changes in maximum CENP-A intensity (Fig. S2D-E). Taken together, these results suggest that ATR’s role in protecting CENP-A occupancy at interphase centromeres is DNA damage-indepdendent and unrelated to its activity as part of the DNA damage response.

### ATR promotes DAXX association with PML NBs

Given our evidence that ATR’s role in protecting CENP-A occupancy at centromeres is DNA damage-independent, we focused our efforts on characterizing basal ATR activity in unperturbed cells. Previous studies have demonstrated that ATR co-localizes with RPA in promyelocytic leukemia nuclear bodies (PML NBs) during interphase and in the absence of DNA damage (Barr et al., 2003). Consistent with this report, we observed robust pRPA2 S33 signal at approximately 30% of PML NBs in U2OS cells (Fig. 3A). The percentage of pRPA2 S33-positive PML NBs was significantly reduced upon acute ATR inhibition (Fig. 3A-B), indicating that ATR is indeed present and active at a subset of PML NBs in unperturbed interphase cells. This led us to hypothesize that ATR activity at PML NBs may indirectly regulate centromere identity and integrity. While ATR was previously shown to localize to mitotic centromeres, ATR is not enriched at interphase centromeres (Kabeche et al., 2018; Fig. S3A), and pRPA2-S33 signal is far more prevalent at PML NBs than at centromeres in unperturbed interphase cells (Fig. S3B). This further demonstrates that ATR’s role in protecting CENP-A occupancy at interphase centromeres may be dependent on its DNA damage-independent activity at PML NBs.

**Figure 3:**
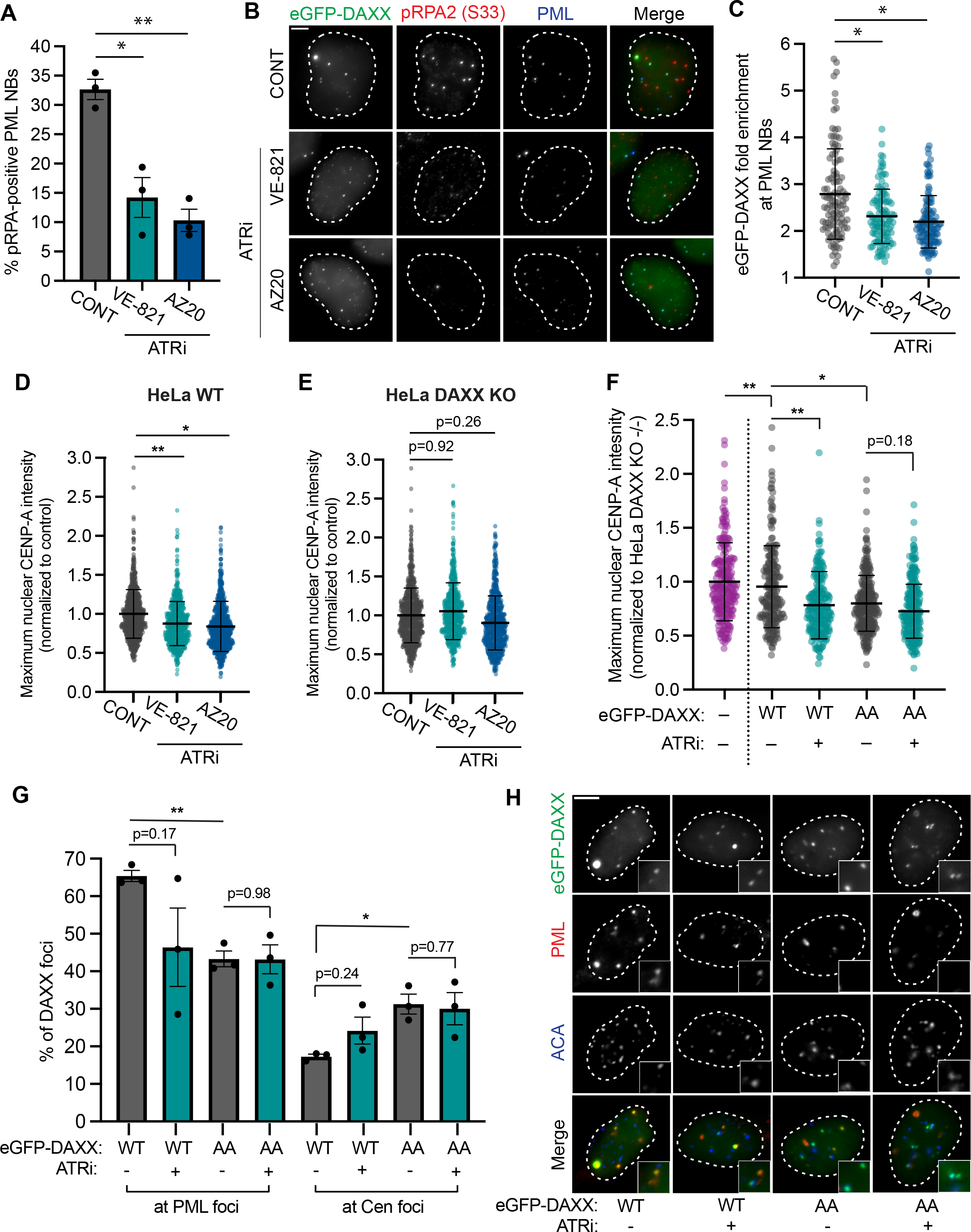
ATR kinase activity and C-terminal DAXX phosphorylation promotes DAXX association with PML NBs. **(A)** *ATR inhibition reduces the proportion of pRPA (S33)-positive PML nuclear bodies*. Asynch. U2OS cells were untreated (CONT) or treated for 1 hour with 10μM VE-821 or 10μM AZ20 (ATRi). (>50 cells per condition; n = 3 biological replicates). Error bars represent mean ±SEM. **(B-C)** *ATR inhibition reduces eGFP-DAXX enrichment at PML nuclear bodies*. (B) Representative images of asynch. U2OS cells transiently transfected with eGFP-DAXX (green). Cells were treated as in (A) and stained for pRPA S33 (red) and PML (blue). Nuclei were identified by DAPI staining and outlined in white. (C) Quantification of eGFP-DAXX fold-enrichment at PML NBs. Mean eGFP-DAXX intensity was quantified at the 3 brightest PML foci and normalized to total nuclear eGFP-DAXX intensity. (>30 cells per condition; n = 3 biological replicates). Outliers were removed according to the ROUT (Q=1%) method. Error bars represent mean ±SD. **(D-E)** *CENP-A loss with ATR inhibition is DAXX-dependent*. Quantification of maximum nuclear CENP-A intensity in asynch. HeLa WT (parental) (D) or HeLa DAXX KO (E) cells treated as in (A). (>130 cells per condition; n = 3 biological replicates). Error bars represent mean ±SD. **(F)** *Phospho-null mutation of eGFP-DAXX decreases maximum nuclear CENP-A intensity and de-sensitizes cells to ATR inhibition*. Asynch. HeLa DAXX KO cells were un-transfected (-) or transiently transfected with eGFP-DAXX^WT^ or eGFP-DAXX^AA^ (S707A;S712A). Cells were untreated (-) or treated for 1 hour with 10μM VE-821 (+ATRi). (>60 cells per condition; n=3 biological replicates). Error bars represent mean ±SD. **(G-H)** *eGFP-DAXX phospho-null mutants are mis-localized to centromeres*. (G) Proportion of eGFP-DAXX foci localizing to PML NBs or centromeres in HeLa DAXX KO cells transiently transfected with eGFP-DAXX^WT^ or eGFP-DAXX^AA^. Cells were untreated (-) or treated for 1 hour with 10μM VE-821 (+ATRi). (>30 cells per condition; n=3 biological replicates). Error bars represent mean ±SEM. (H) Representative images of eGFP-DAXX-expressing cells quantified in (G). Cells were stained for PML (red) and anti-centromere antibody (ACA; blue). Nuclei were identified by DAPI staining and outlined in white. Inset depicts zoomed-in image of 2 eGFP-DAXX foci for better visualization of DAXX localization. *p ≤ 0.05, **p ≤ 0.01, two-tailed t-test of replicate averages. Scale bars for all panels = 5 μm.

To determine how ATR activity at PML NBs may regulate centromere identity, we focused on PML NB components with dual roles at centromeric chromatin. DAXX and ATRX are commonly associated with PML NBs, but these histone H3.3 chaperones also localize to centromeres to regulate heterochromatin formation and centromere transcription (Morozov et al., 2012; Morozov et al., 2017). However, U2OS cells do not have functional ATRX (Heaphy et al., 2011) but nonetheless display CENP-A loss from centromeres when treated with ATR inhibitors (Fig. 1). Furthermore, DAXX is specifically implicated in CENP-A mis-localization from centromeres (Nye et al., 2018; Shrestha et al., 2021), and phosphorylation of DAXX at residues matching ATR’s consensus motif has been detected in previous studies (Matsuoka et al., 2007; Stokes et al., 2007). This led us to hypothesize that basal ATR activity may protect CENP-A occupancy at centromeres by promoting the sequestration of DAXX in PML NBs.

To test this hypothesis, we expressed eGFP-DAXX in U2OS cells. eGFP-DAXX strongly co-localized with PML NBs and was present at all pRPA2 S33-positive PML foci (Fig. 3B). By measuring fold enrichment of eGFP signal at the three brightest PML foci relative to total nuclear signal, we found that treatment with VE-821 and AZ20 for 1 hour resulted in diffuse GFP signal throughout the nucleus (Fig. 3B) and reduced enrichment at PML NBs by approximately 20% (Fig. 3C). We also measured the percentage of PML NBs that co-localized with eGFP-DAXX foci and saw a similar, albeit less significant decrease in DAXX-positive PML NBs (Fig. S3C). This demonstrates that ATR may promote DAXX sequestration in PML NBs.

Next, we sought to determine whether CENP-A loss from interphase centromeres with ATR inhibition is DAXX-dependent. To test this, we obtained commercially available HeLa DAXX-knockout cells (HeLa DAXX KO [Abcam]). We first confirmed that treatment of HeLa DAXX KO cells with VE-821 and AZ20 reduced ATR kinase activity as measured by western blotting for pRPA2 S33 (Fig. S3D). We also determined that whole-cell CENP-A levels were unchanged by DAXX knock-out in comparison to the parental HeLa cell line (HeLa WT [Abcam]; Fig. S3D). While maximum CENP-A immunofluorescence intensity was decreased in HeLa WT cells treated with ATR inhibitors (Fig. 3D), no significant changes in CENP-A intensity were detected in HeLa DAXX KO cells (Fig. 3E). This demonstrates that CENP-A loss with ATR inhibition is indeed DAXX-dependent.

Given that ATR inhibition decreases DAXX association with PML NBs and leads to a DAXX-dependent decrease in CENP-A occupancy at interphase centromeres, we also expected to see an increase in DAXX localization to interphase centromeres. To assess this, we performed ChIP of endogenous DAXX to determine changes in centromere occupancy following ATR inhibition. In asynchronous U2OS cells, we saw a general increase in DAXX localization at interphase centromeres using two distinct α-satellite-specific primer pairs (Fig S3E). However, the magnitude of this increase varied significantly across biological replicates, suggesting that DAXX may only localize to a subset of interphase centromeres, or that DAXX association with centromeres is transient following ATR inhibition. Consistent with DAXX’s general increase at interphase centromeres, ATR inhibition also resulted in an increase in histone H3.3 occupancy at interphase centromeres (Fig. S3F), suggesting that DAXX localization to centromeres coincides with the deposition of H3.3-containting nucleosomes. This increase in centromeric H3.3 occupancy may explain why histone H2A levels remained unchanged (Fig. S1D) despite a consistent decrease in CENP-A nucleosome occupancy with ATR inhibition (Fig. 1). Taken together, these findings demonstrate that ATR promotes DAXX association with PML NBs, which may limit DAXX localization to centromeres as well as aberrant H3.3 deposition and CENP-A eviction.

### Mutation of C-terminal DAXX residues phenocopies ATR inhibition

ATR modulates the activity of its targets through phosphorylation of serine or threonine residues preceding a glutamine (S/T-Q) (Kim et al., 1999). Previous studies have identified phosphorylation of DAXX at ATR consensus motifs in response to genotoxic stress (Matsuoka et al., 2007; Stokes et al., 2007), but this has not been assessed in unperturbed cells. DAXX association with PML NBs is mediated through their respective SUMOylated residues and SUMO-interacting motifs (SIMs) (Lin et al., 2006). Phosphorylation of DAXX in or near its C-terminal SIM has been shown to modulate its affinity for SUMOylated residues on PML protein, thereby regulating DAXX-PML association (Chang et al., 2011). This suggests that ATR may directly phosphorylate DAXX near its SIM to regulate DAXX localization. To assess this possibility, we mapped previously-identified S/T-Q phosphorylation of DAXX protein (PhosphoSitePlus), which led to the identification of S707 and S712 as putative ATR substrates residing near the C-terminal SIM of DAXX(Chang et al., 2011; Matsuoka et al., 2007; Stokes et al., 2007). To assess whether phosphorylation of these sites impacts CENP-A occupancy, we generated eGFP-DAXX phospho-null double mutants in which these serine residues were replaced by alanine residues (S707A;S712A, “AA”). We expressed eGFP-DAXX^WT^ or eGFP-DAXX^AA^ in HeLa DAXX KO cells and measured maximum nuclear CENP-A intensity via immunofluorescence microscopy following ATR inhibition (Fig. 3F, Fig. S3H). Similar to HeLa DAXX WT cells, treatment of HeLa DAXX KO cells expressing eGFP-DAXX^WT^ with VE-821 for 1 hour resulted in a strong decrease in maximum CENP-A intensity. Surprisingly, expression of eGFP-DAXX^AA^ mimicked this decrease, and treatment of these cells with VE-821 had no significant effect on CENP-A occupancy. This suggests that S707 and S712 of DAXX may be bona fide ATR substrate, and that phosphorylation of these sites may protect CENP-A occupancy at interphase centromeres.

Next, we assessed whether eGFP-DAXX^AA^ displayed defects in PML NB and centromere localization. We measured the percentage of eGFP-DAXX^AA^ or eGFP-DAXX^WT^ foci localized to PML NBs or centromeres. In untreated cells, approximately 65% of eGFP-DAXX^WT^ foci co-localized with PML NBs, while a small subset (∼15%) co-localized with centromeres (Fig. 3G-H). As expected, we saw a decrease in the percentage of eGFP-DAXX^WT^ foci at PML NBs, which correlated with an increase in centromere localization upon treatment with VE-821 for 1 hour. While these changes were not statistically significant, these complimentary defects in localization demonstrate that translocation of DAXX from PML NBs to centromeres may be occurring upon ATR inhibition. Moreover, eGFP-DAXX^AA^ foci displayed a similar localization pattern to that of eGFP-DAXX^WT^ in ATR-inhibited cells, in which approximately 45% of eGFP-DAXX^AA^ co-localized with PML NBs. However, these mutants displayed strong centromere localization, with approximately 30% of eGFP-DAXX^AA^ foci localizing to centromeres. Treatment with VE-821 had no effect on the localization pattern of these eGFP-DAXX^AA^ mutants, demonstrating that these phospho-null mutations confer resistance to the effects of ATR inhibition on DAXX localization. Taken together, these findings provide strong evidence that phosphorylation of S707 and S712 by ATR may regulate DAXX association with PML NBs, thereby limiting DAXX localization to centromeres to protect CENP-A occupancy.

### CENP-A loss from centromeres persists until G1 of the following cell cycle, leading to errors in mitotic chromosome segregation

Long-term inhibition or loss of ATR can lead to mitotic catastrophe and chromosome mis-segregation due to the accumulation of DNA damage in interphase (Brown and Baltimore, 2000), and acute ATR inhibition in mitosis results in chromosome mis-segregation due to impaired Aurora B signaling at centromeres (Kabeche et al., 2018). However, it is unknown whether acute ATR inhibition during interphase impacts mitotic fidelity. Proper CENP-A occupancy at interphase centromeres is crucial to build a functional kinetochore in mitosis (Régnier et al., 2005). Given that acute ATR inhibition reduces CENP-A occupancy in interphase, we asked whether CENP-A loss from centromeres persists through mitosis after ATR activity has been restored. We treated U2OS cells with VE-821 for 1 hour, then collected cells for immunofluorescence microscopy and western blotting at various time points following inhibitor wash-out (Fig. 4A). ATR activity, as measured by pRPA2 S33 signal, was fully restored within 1 hour after inhibitor wash-out. In contrast, CENP-A intensity remained diminished 1 hour and 6 hours after ATR activity was restored (Fig. 4B). Given that CENP-A deposition only occurs in early G1 (Jansen et al., 2007), we hypothesized that CENP-A may not be replenished at centromeres until the following cell cycle. Similar to our previous experiment, we treated U2OS cells with VE-821 for 1 hour, after which we washed out ATR inhibitor. We then treated cells with Nocodazole for 16 hours to synchronize cells in prometaphase and released cells into G1. We confirmed that our initial ATR inhibition was successful and that cells collected for immunofluorescence microscopy were indeed in G1 by western blotting for pRPA2 S33 and cyclin A, respectively (Fig. S4A). Further, we determined that G1 cells did not have drastically increased levels of DNA damage compared to asynchronous control cells (Fig. S4A). We found that maximum CENP-A intensity of G1 cells treated with VE-821 in the previous cell cycle was identical to that of control cells (Fig. 4C), demonstrating that CENP-A occupancy at centromeres is indeed restored in G1.

**Figure 4:**
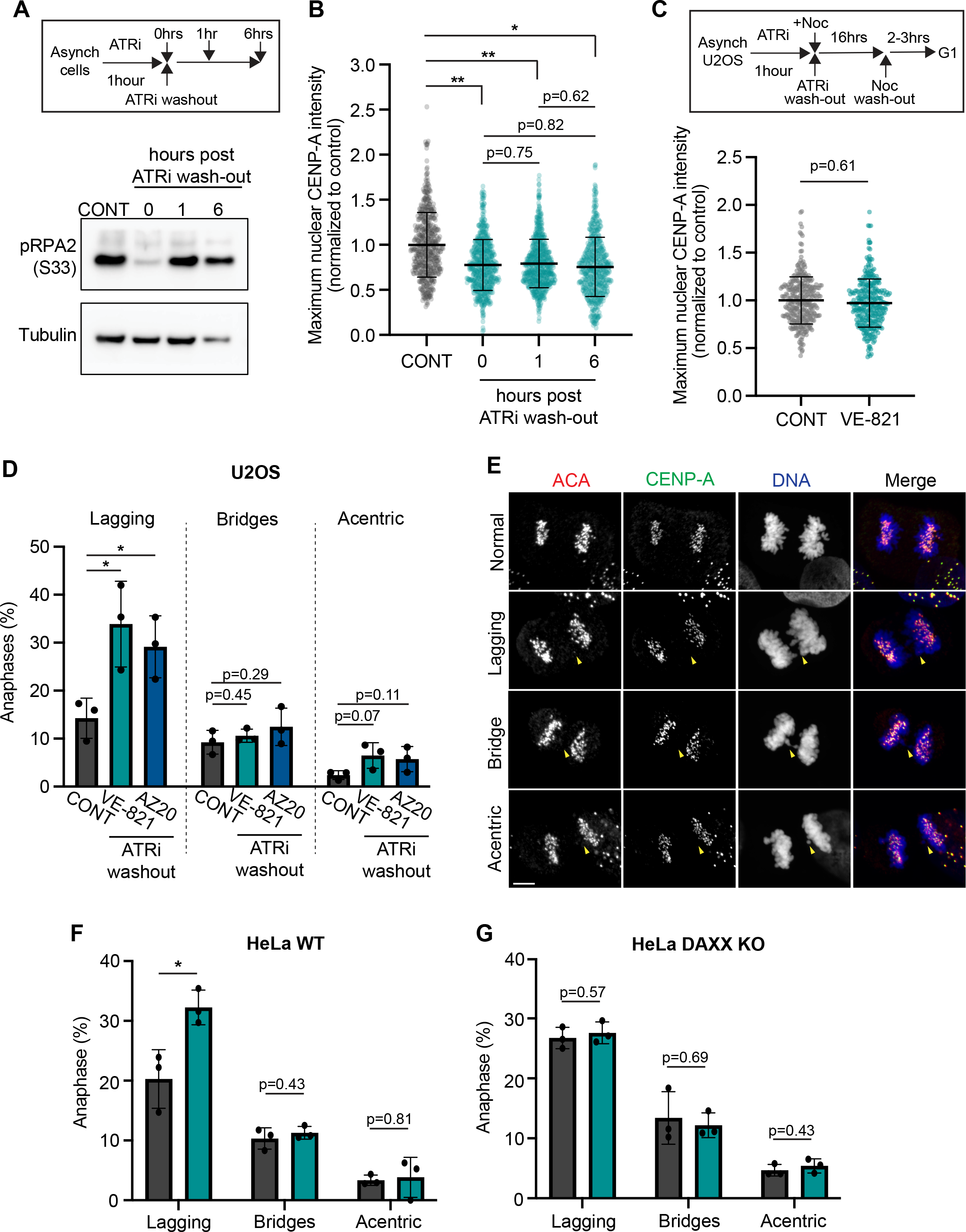
CENP-A loss from centromeres persists until G1 of the following cell cycle, leading to errors in mitotic chromosome segregation. **(A)** *ATR activity is restored following inhibitor wash-out*. (Top) Schematic of ATR inhibitor (ATRi) wash-out experiments. Asynchronous cells were untreated (CONT) or treated for 1 hour with 10μM VE-821 or 10μM AZ20, after which ATR inhibitors were washed out. Cells were collected for western blotting and immunofluorescence microscopy just after wash-out (0 hours) or at 1 and 6 hours post wash-out. (Bottom) Representative western blot of pRPA (S33) levels in ATRi wash-out experiments. **(B)** *CENP-A intensity remains depleted following ATR inhibitor wash-out*. Quantification of maximum nuclear CENP-A intensity in asynchronous (asynch). U2OS treated with 10μM VE-821 and prepared as described in (A). (>150 cells per condition; n = 3 replicates). Outliers were removed according to the ROUT (Q=1%) method. Error bars represent mean ±SD. **(C)** *CENP-A levels are not restored until G1 of the following cell cycle*. (Top) Schematic of ATRi and Nocodazole (Noc) wash-out experiments for isolation of G1 cells. Asynchronous cells were untreated (CONT) or treated for 1 hour with 10μM VE-821, after which VE-821 was washed out. Cells were then incubated with Nocodazole (Noc; 50 ng/mL) for 16 hours, after which Noc was washed out and G1 cells were collected for western blotting and immunofluorescence microscopy. (Bottom) Quantification of maximum nuclear CENP-A intensity in G1 U2OS cells (>90 cells per condition; n = 3 replicates). Outliers were removed according to the ROUT (Q=1%) method. Error bars represent mean ±SD. **(D-E)** *Acute ATR inhibition in interphase increases the incidence of lagging chromosomes in mitosis*. (D) Quantification of anaphases with lagging chromosomes (lagging), chromosome bridges (bridges) or acentric fragments (acentric) in asynch. U2OS cells 6 hours after ATRi wash-out as described in (A). (>100 cells per condition; n=3 biological replicates). Error bars represent mean ±SEM. (E) Representative images of anaphase cells quantified in (D). Cells were stained for anti-centromere antibody (red), CENP-A (green), and DNA (DAPI; blue). Yellow arrows indicate lagging chromosomes. Scale bar = 5 μm. (**F-G**) *Chromosome segregation defects arising from acute ATR inhibition during interphase is DAXX-dependent*. Quantification of anaphases with lagging chromosomes (lagging), chromosome bridges (bridges) or acentric fragments (acentric) in asynch. HeLa WT (parental) cells (F) or HeLa DAXX KO cells (G) 6 hours after 10μM VE-821 wash-out as described in (A). (>30 cells per condition; n=3 biological replicates). Error bars represent mean ±SEM. *p ≤ 0.05, **p ≤ 0.01, two-tailed t-test of replicate averages.

Given that CENP-A loss persists after ATR activity is restored, we asked how decreased CENP-A occupancy at interphase centromeres affects mitotic chromosome segregation. Because ATR has a crucial role in promoting mitotic kinetochore-microtubule error correction (Kabeche et al., 2018), we repeated our wash-out experiment to isolate the effects of CENP-A loss due to acute ATR inhibition in interphase (Fig. 4A). U2OS cells were treated with VE-821 or AZ20 for 1 hour, and mitotic anaphase cells were identified 6 hours after inhibitor wash-out. Surprisingly, these cells exhibited a 2-fold increase in the incidence of lagging chromosomes and a moderate but non-significant increase in acentric fragments, but an increase in chromosome bridges was not detected (Fig. 4D). Interestingly, many of the lagging chromosomes we observed possessed anti-centromere antibody signal, but often did not contain CENP-A signal (Fig. 4E), suggesting that these mis-segregation events may be due to CENP-A loss.

Finally, we sought to determine if the increase in the incidence of lagging chromosomes after acute ATR inhibition in interphase was DAXX-dependent. We repeated our inhibitor wash-out experiments in HeLa DAXX KO and HeLa WT cells and assessed mitotic anaphases 6 hours after VE-821 wash-out. We confirmed that ATR activity is restored after VE-821 wash-out in both cell lines (Fig. S4B-C). As with U2OS cells, we detected an increase in lagging chromosomes in HeLa WT cells transiently treated with VE-821 (Fig. 4F). However, the incidence of lagging chromosome in HeLa DAXX KO cells was unaffected by transient ATR inhibition (Fig. 4G). These data strongly support that acute ATR inhibition causes a DAXX-dependent loss of CENP-A from centromeres, resulting in mitotic chromosome segregation defects.

## Discussion

Our results show that basal ATR activity in the absence of DNA damage protects CENP-A nucleosome occupancy at centromeres by promoting DAXX association with PML NBs. While DAXX localization to centromeres is increased upon ATR inhibition, we also observed a similar increase in DAXX localization to other sites in which H3.3 nucleosomes are regularly deposited, including peri-centromeres and telomeres (Fig. S4D-E). This suggests that ATR inhibition may induce a general “release” of DAXX from PML NBs, rather than targeted translocation to centromeres. Thus, DAXX-dependent CENP-A loss from interphase centromeres is an important consequence of ATR inhibition, but activity at other genomic loci is likely to be increased as well.

Given that DAXX-dependent H3.3 deposition at these loci modulates transcription and heterochromatin formation (Ahmad and Henikoff, 2002), an increase in DAXX localization upon ATR inhibition may result in widespread chromatin and transcriptomic changes. Whether or not these changes occur in response to genotoxic stress as well as acute ATR inhibition is a question worth considering.

We also showed that CENP-A loss from interphase centromeres with acute ATR inhibition persists after ATR activity is restored, and that CENP-A occupancy is not replenished until G1 of the following cell cycle. This sustained CENP-A loss correlates with an increase in lagging chromosomes during anaphase (Fig. 4). It remains unclear how CENP-A loss alone could increase the incidence of lagging chromosomes. It is possible that aberrant H3.3 deposition at interphase centromeres leads to changes in centromeric chromatin structure that alter the recruitment of kinetochore components. Alternatively, eviction of H3.3 nucleosomes after ATR activity is restored may lead to a net loss of centromeric nucleosomes. Thus, multiple mechanisms may contribute to an increase in mitotic chromosome segregation errors with acute ATR inhibition, highlighting the need to better understand how nucleosome dynamics affect centromere and kinetochore structure.

While the full implications of this pathway remain an exciting question, our work demonstrates that partial loss of CENP-A from centromeres with acute ATR inhibition can lead to genome instability via increased chromosome segregation defects. ATR inhibitors are currently used as cancer therapeutics, but dose-limiting toxicities have been observed in some cases (Barnieh et al., 2021). Given that short-term ATR inhibition is sufficient to produce detrimental mitotic defects, pulsatile administration of ATR inhibitors may help to overcome these challenges while preserving therapeutic efficacy. Furthermore, our results demonstrate that CENP-A loss and chromosome mis-segregation with acute ATR inhibition is DAXX-dependent, suggesting that ATR inhibitors may be more potent in DAXX-proficient cells. Thus, DAXX may be a suitable biomarker for ATR inhibitor efficacy. We hope that these findings will motivate future studies to fully elucidate this pathway and uncover new therapeutic paradigms for ATR inhibitors.

## Acknowledgement

We thank Drs. Xiaolu Yang, Torsten Wittman, and Bob Weinberg for reagents, and members of the Yale Cancer Institute for discussions. This work is supported by the Pershing Square Sohn Award.

## Author contributions

I.T and L.K. designed the study. I.T., E.B, Y.K. and L.K. performed the experiments and analyses. L.K. supervised the experiments and analyses. I.T and L.K. prepared the manuscript with contributions from all authors.

## Competing interests

All authors do not have competing interests.

## STAR * METHODS

### KEY RESOURCES TABLE

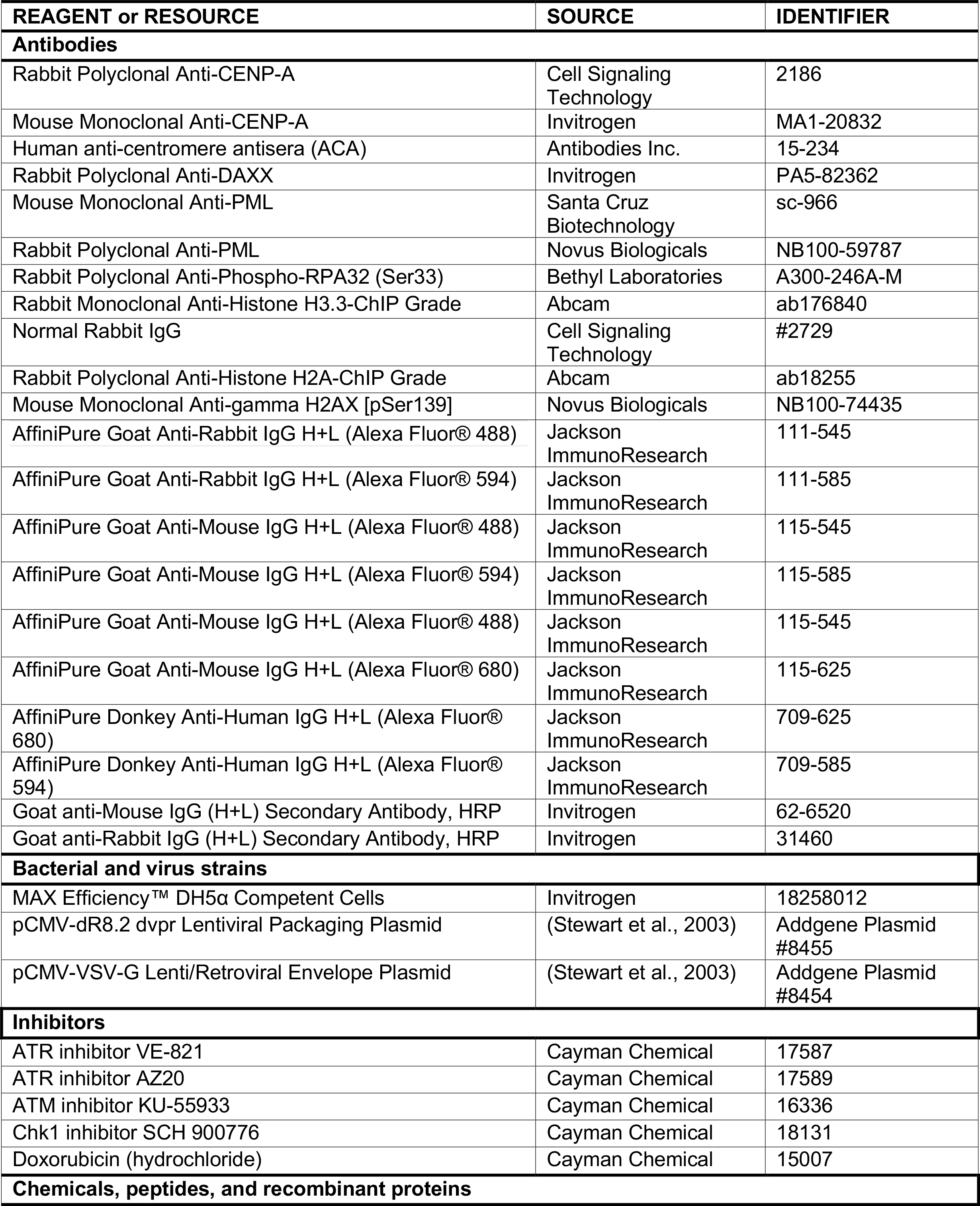
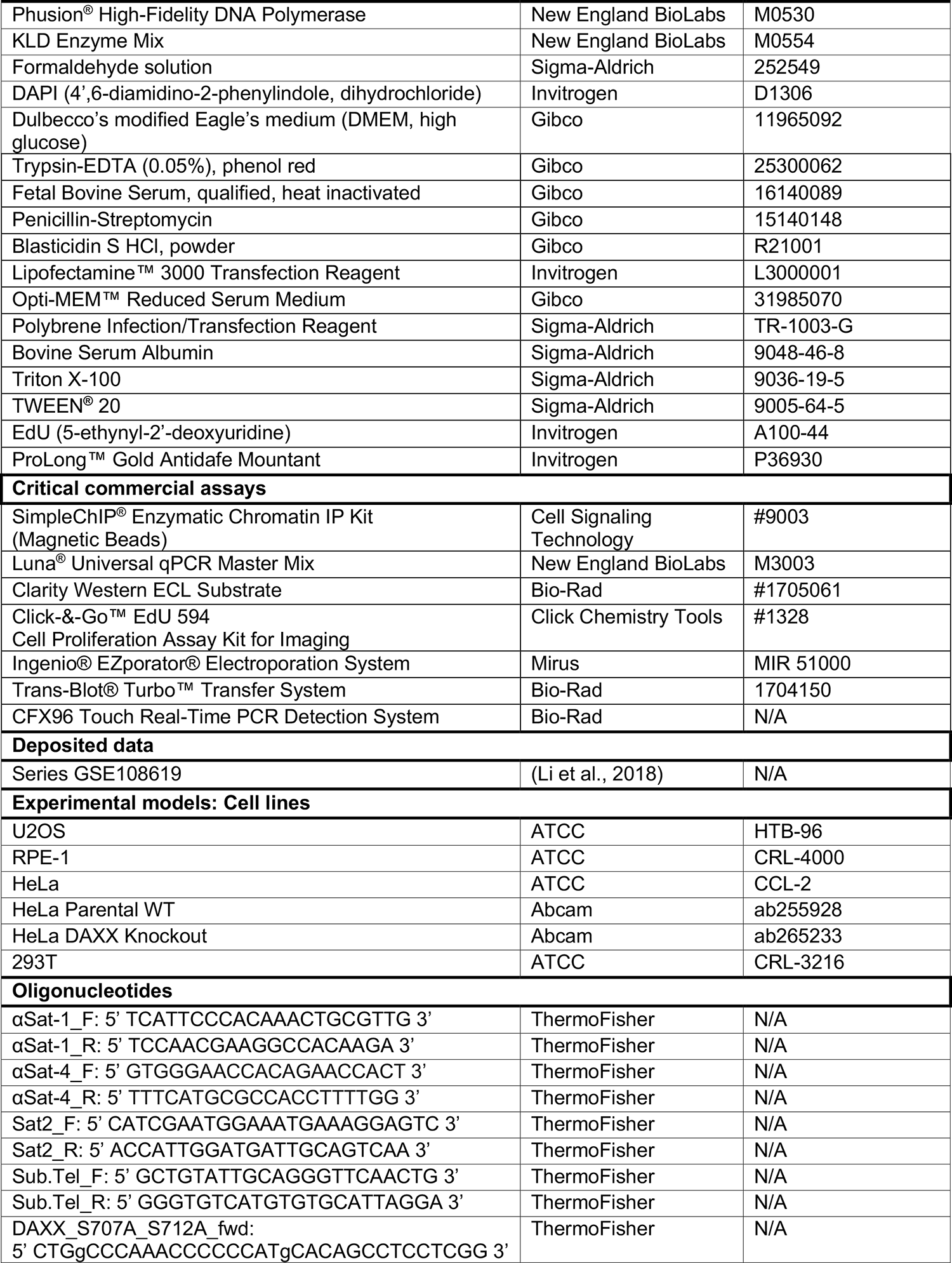
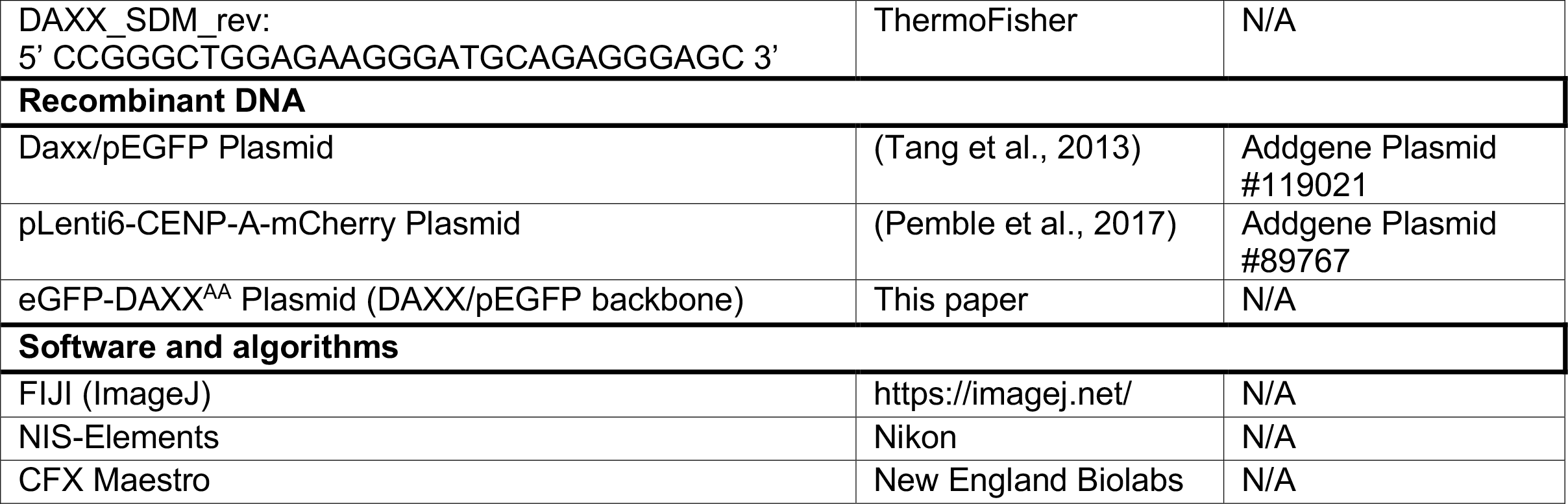

### RESOURCE AVAILABILITY

#### Lead Contact

Further information and requests for reagents should be direct to and will be fulfilled by the lead contact Lilian Kabeche.

#### Materials Availability

Requests for cell lines and plasmids generated in this study should be directed to the lead contact

### EXPERIMENTAL MODEL AND SUBJECT DETAILS

#### Cell lines and culture medium

U2OS (ATCC, HTB-96), HeLa (ATCC, CCL-2), RPE-1 (ATCC, CRL-4000), HeLa WT (Abcam, ab255928), HeLa DAXX Knockout (Abcam, ab265233), and 293T (ATCC, CRL-3216) cells were cultured in Dulbecco’s modified Eagle’s medium (DMEM; Gibco) supplemented with 10% fetal bovine serum (FBS; Gibco) and 1% penicillin/streptomycin (Gibco). Media for U2OS cells stably expressing CENP-A-mCherry was supplemented with 10 µg/mL Blasticidin S HCl (Gibco) for cell selection and maintenance.

### METHOD DETAILS

#### Cell transfection

U2OS and HeLa DAXX Knockout cells were transiently transfected with eGFP-DAXX^WT^ plasmid (DAXX/pEGFP, gift from Xiaolu Yang, Addgene 119021) or eGFP-DAXX^AA^ plasmid using the Ingenio® EZporator® Electroporation System (Mirus) according to manufacturer instructions. Transfected cells were analyzed the following day.

#### Viral Infection

CENP-A-mCherry lentivirus was generated via co-transfection of 293T cells with pLenti6-mCherry-CENP-A plasmid (gift from Torsten Wittmann, Addgene 89767), pCMV dR8.2 dvpr (Addgene 8455, gift from Bob Weinberg), and pCMV-VSV-G (Addgene 8454, gift from Bob Weinberg) using Lipofectamine™ 3000 Transfection Reagent (Invitrogen) according to manufacturer instructions. Transfected 293T cells were cultured in reduced serum medium (Opti-MEM; Invitrogen) without penicillin/streptomycin, then split into Dulbecco’s modified Eagle’s medium (DMEM; Gibco) supplemented with 10% fetal bovine serum (FBS; Gibco) the following day. Viral supernatant was harvested 2 days after transfection and added to U2OS cells cultured in penicillin/streptomycin-free DMEM supplemented with 10% FBS and 8µg/mL polybrene infection/transfection reagent (Sigma-Aldrich). U2OS cells were analyzed for successful expression of CENP-A-mCherry 2 days after infection and selected using 10 µg/mL Blasticidin S HCl (Gibco).

#### Site-directed mutagenesis

Serines 707 and 712 of eGFP-DAXX (DAXX/pEGFP, gift from Xiaolu Yang, Addgene 119021) were mutated to alanine residues via site-directed mutagenesis. Oligonucleotide primers introducing these point mutations were obtained from ThermoFisher (DAXX_S707A_S712A_fwd: 5’ CTGgCCCAAACCCCCCATgCACAGCCTCCTCGG 3’ and DAXX_SDM_rev: 5’ CCGGGCTGGAGAAGGGATGCAGAGGGAGC 3’) and used to amplify eGFP-DAXX via PCR using Phusion^®^ High-Fidelity DNA Polymerase (New England BioLabs). Phosphorylation, ligation, and template removal were performed using KLD Enzyme Mix (New England BioLabs) according to manufacturer instructions. The reaction was then transformed into MAX Efficiency™ DH5αCompetent Cells (Invitrogen). Plasmid DNA from bacterial cultures was prepped according to standard Alkaline Lysis Miniprep protocols and sequenced through GENEWIZ (Azenta Life Sciences).

#### Western Blotting

Cells were cultured in 6-well tissue culture plates. Confluent cells were incubated with 500 uL of 0.05% Trypsin-EDTA (Gibco) at 37ºC for 10 minutes and collected in an Eppendorf tube. Trypsinized cells were spun down for 1 minute at 13,000 rpm in a centrifuge, then resuspended in 25 uL PBS and 25 uL of 2x SDS Sample Buffer. The protein samples were lysed with an insulin syringe and incubated at 95ºC for 15 minutes. Protein samples were run on a 7.5% SDS-PAGE gel at 100V until optimal separation was achieved. Protein gels were transferred to 0.2 µm nitrocellulose membranes using the Trans-Blot® Turbo™ Transfer System (Bio-Rad). Membranes were stained for 5 min with Ponceau S staining solution (0.5% (w/v) Ponceau S, 1% glacial acetic acid) to ensure equal protein loading and then washed with 1X Tris-Buffered Saline + 0.1% Tween 20 detergent (TBS-T). Membranes were blocked with TBS-T + 5% bovine serum albumin (BSA) or milk for 1 hour at room temperature. Primary antibodies were diluted 1:1000 in blocking solution, then added to membranes for overnight incubation at 4ºC. Membranes were then washed 3 times with TBS-T for 5 minutes. Secondary horseradish peroxidase (HRP)-conjugated secondary antibodies (Invitrogen) were diluted 1:1000 in blocking solution and added to membranes for 1 hour at room temperature, followed by an additional 3 washes. Membranes were incubated for 5 min with Clarity Western ECL substrate (Bio-Rad) and imaged on the ChemiDoc MP Imaging system (Bio-Rad).

#### Chromatin Immunoprecipitation and quantitative PCR

Chromatin immunoprecipitation was carried out per the SimpleChIP^®^ Enzymatic Chromatin IP Kit with Magnetic Beads (Cell Signaling Technology). Briefly, cells were cultured in 15 cm flasks with a minimum of 2 confluent flasks per condition. After treatment with ATR inhibitors for 1 hour, cells were incubated with 0.05% Trypsin-EDTA (Gibco) at 37ºC for 10 minutes. Trypsinized cells were collected in 50 mL conical tubes and resuspended in fresh media. Cross-linking was carried out by addition of 37% formaldehyde solution (Sigma-Aldrich, final concentration of 1%), followed by quenching with glycine. Cells were then washed with ice-cold PBS. Nuclei Preparation and Chromatin Digestion was carried out using the buffers and micrococcal nuclease provided by the SimpleChIP^®^ kit according to manufacturer instructions. Cell lysis was performed using insulin syringes. Optimal chromatin digestion was analyzed via DNA gel electrophoresis to ensure that DNA was digested to a length of approximately 150-900 bp. Chromatin immunoprecipitation was carried out according to manufacturer instructions using approximately 5 ug of digested, cross-linked and 1 µg of Rabbit Polyclonal Anti-CENP-A antibody (Cell Signaling Technology), Rabbit Polyclonal Anti-Histone H2A antibody (Abcam), Rabbit Polyclonal Anti-DAXX antibody (Invitrogen), Rabbit Monoclonal Anti-Histone H3.3 antibody (Abcam), or Normal Rabbit IgG (Cell Signaling Technology). Chromatin elution, reversal of cross-links, and DNA purification using spin columns was carried out using the reagents and instructions provided in the SimpleChIP^®^ kit. Quantitative PCR reactions were prepared using the Luna^®^ Universal qPCR Master Mix (New England BioLabs) in technical triplicate to amplify centromere, peri-centromere, or telomere DNA and run on the CFX96 Touch Real-Time PCR Detection System (Bio-Rad). C_q_ values were calculated for each SYBR fluorescence trace using the CFX Maestro software. Percentage of input was calculated according to the Delta C_q_ values for each sample.

#### Immunofluorescence microscopy

Cells incubated on coverslips were fixed with 3.5% paraformaldehyde for 15 minutes, permeabilized with Phosphate Buffered Saline (PBS) + 0.5% Triton X-100 for 10 minutes, and blocked with PBS + 1% bovine serum albumin (BSA) for 30 minutes. For cells that were treated with 5-ethynyl-2’-deoxyuridine (EdU), the Click-&-Go EdU Cell Proliferation Assay Kit (Click Chemistry Tools) was used to label newly synthesized DNA with Azide 594. This reaction was performed according to manufacturer instructions prior to incubation with primary antibodies. Coverslips were washed with PBS for 5 minutes after EdU labeling. Cells on coverslips were then incubated with primary antibody diluted in PBS + 1% BSA at 4°C overnight. Cells were then washed with PBS for 5 minutes. Secondary antibodies conjugated to fluorophores were diluted in PBS + 1% BSA. Cells on coverslips were then incubated with secondary antibodies and 1µg/mL DAPI stain for 1 hour at room temperature (∼25°C). Cells were washed again with PBS for 5 min and mounted onto glass slides using ProLong™ Gold Antifade Mountant (Invitrogen) and left to dry overnight. All images were acquired on a Nikon ECLIPSE Ti2 widefield epi-fluorescence microscope with a Hamamatsu Fusion sCMOS camera. Image acquisition was managed using NIS-Elements (Nikon). All images were analyzed in FIJI (ImageJ) and processed in Adobe Photoshop.

#### Live-cell imaging and quantitative analysis of U2OS CENP-A-mCherry immunofluorescence intensity

Asynchronous U2OS cells stably expressing CENP-A-mCherry were cultured in a 4-chamber glass-bottom dish (Cellvis). The dish was mounted on a Tokai Hit STX stage-top incubator (Spectra Services) to maintain an imaging environment of 37ºC with 5% CO2. Coordinates were set for interphase U2OS cells expressing CENP-A-mCherry in 3 chambers. Z-stacks were taken at 60x for each cell with a range of 4µm at 0.3 µm steps. After the first set of images (time = 0 mins) were taken, the media in each chamber was immediately replaced with fresh media or media containing 10µM VE-821 or 10µM AZ20. The remaining images were taken at 10-minute intervals over the course of 1 hour. Images were analyzed in FIJI (ImageJ). Average intensity projections were compiled for each z-stack at each time point. For each cell at time = 0 mins, the three brightest CENP-A-mCherry foci were identified using the *Find Maxima* function. A circle with a diameter of 0.2μm was drawn for each CENP-A-mCherry maximum and measured for each time point. Maxima values were then normalized to the total nuclear intensity for that cell and time point. The normalized values in each cell for all time points were divided by the value at time = 0 mins to display maximum CENP-A-mCherry intensity with 100% at time = 0 mins.

#### Quantitative analysis of maximum CENP-A immunofluorescence intensity

For quantitative analysis of CENP-A intensity, unsynchronized cells were cultured on coverslips and incubated with various inhibitors for 1 hour. Coverslips were prepared for immunofluorescence microscopy as described above and immuno-stained using Rabbit Polyclonal Anti-CENP-A antibody (Cell Signaling Technology) and Goat Anti-Rabbit antibody conjugated to Alexa Fluor 488 (Jackson InmmunoResearch). Single-stack images were taken at 60x and analyzed in FIJI (ImageJ). Interphase nuclei were masked according to DAPI staining, and maximum nuclear CENP-A fluorescence intensity was quantified for all conditions. Mean background values from each image were subtracted from all maximum nuclear CENP-A intensity values corresponding to that image. In order to display the results from each biological replicate in one graph, all intensity values were normalized to the average control value for that image set, resulting in a mean intensity value of 1.0 for control cells.

#### Cell cycle sorting via immunofluorescence microscopy

To sort cells into their distinct cell cycle phases, unsynchronized cells were cultured on coverslips and incubated with ATR inhibitors for 1 hour and 10 µM 5-ethynyl-2’-deoxyuridine (EdU) for 30 minutes. EdU was labeled using the Click-&-Go™ EdU 594 Cell Proliferation Assay Kit for Imaging click chemistry (Click Chemistry Tools) and cells were stained for CENP-A as described above. Single-stack images were taken at 60x and analyzed in FIJI (ImageJ). Interphase nuclei were masked according to DAPI staining. The mean nuclear DAPI and EdU-594 fluorescence intensity as well as nuclear area were measured for each cell in all conditions. Mean background values for both DAPI and EdU-594 channels from each image were subtracted from all mean DAPI and EdU-594 intensity values corresponding to that image. Integrated density of DAPI and EdU-594 signal for each cell was calculated by multiplying mean nuclear intensity (adjusted for background) by nuclear area. EdU-594 integrated density for at least 100 cells was converted to a logarithmic scale and plotted against DAPI integrated density in GraphPad Prism. Cells were assigned to their distinct cell cycle phases according to clustering patterns on the graph. Maximum CENP-A intensity was analyzed as described above for the G1, S, and G2-phase cells identified in that image set. In order to display the results from each biological replicate in one graph, all intensity values were normalized to the average control value for that image set, resulting in a mean intensity value of 1.0 for control cells.

#### Quantitative analysis of mean γH2AX immunofluorescence intensity

For quantitative analysis of γH2AX intensity, unsynchronized cells were cultured on coverslips and incubated with various inhibitors for 1 hour. Coverslips were prepared for immunofluorescence microscopy as described above and immuno-stained using Mouse Monoclonal Anti-gamma H2AX (pSer139) primary antibody (Novus Biologicals) and Goat anti-Mouse secondary antibody conjugated to Alexa Fluor 594 (Jackson ImmunoResearch). Centromeres were marked using anti-centromere primary antibody (ACA; Antibodies Inc.) and Donkey anti-Human secondary antibody conjugated to Alexa Fluor 680 (Jackson ImmunoResearch). Single-stack images were taken at 60x and analyzed in FIJI (ImageJ). Interphase nuclei were masked according to DAPI staining and mean nuclear γH2AX intensity was quantified for all conditions. To measure mean γH2AX foci at centromeres, ACA foci were detected with FIJI’s *Find Maxima* function using an appropriate threshold that remained the same for all conditions. Each ACA focus was enlarged to a circle with a diameter of 0.2μm, and mean γH2AX intensity was measured at each focus and averaged for each cell. Mean background values from each image were subtracted from all γH2AX intensity values corresponding to that image. In order to display the results from each biological replicate in one graph, all intensity values were normalized to the average control value for that image set, resulting in a mean intensity value of 1.0 for control cells.

#### Quantitative analysis of chromosome mis-segregation events

Unsynchronized cells were cultured on coverslips and incubated with ATR inhibitors for 1 hour. Coverslips were prepared for immunofluorescence microscopy as described above and immuno-stained using human anti-centromere antibody (ACA; Antibodies Inc.) to mark centromeres/kinetochores and Mouse Monoclonal Anti-CENP-A primary antibody (Invitrogen). Goat Anti-Mouse secondary antibody conjugated to Alexa Fluor 488 (Jackson ImmunoResearch) was used to label CENP-A, and Donkey Anti-Human secondary antibody conjugated to Alexa Fluor 594 was used to label centromeres (Jackson ImmunoResearch). Representative images were taken at 60x. Lagging chromosomes were identified in anaphase cells if they met the following criteria: (1) the distance between the anaphase planes is ≥8µm, (2) a single ACA focus that localizes to a chromosome (DAPI) can be seen in between the two planes, and (3) this ACA focus is ≥0.5 µm from the closest anaphase plane. Chromosome bridges were identified as elongated DNA fragments stained by DAPI that spanned the length between the two anaphase planes. Acentric fragments were identified as small DNA fragments between two anaphase planes with no ACA staining.

#### Quantitative analysis of %pRPA S33 or DAXX-positive PML nuclear bodies

Asynchronous U2OS cells were transiently transfected with eGFP-DAXX plasmid as described above and cultured on coverslips. The next day, cells were incubated with ATR inhibitors for 1 hour, after which coverslips were prepared for immunofluorescence microscopy as described above. Cells were immuno-stained with Rabbit Polyclonal Anti-Phospho-RPA32 (Ser33) primary antibody (Bethyl Laboratories), Mouse Monoclonal Anti-PML primary antibody (Santa Cruz Biotechnology), Goat anti-Rabbit secondary antibody conjugated to Alexa Fluor 594 (Jackson ImmunoResearch), and Goat Anti-Mouse secondary antibody conjugated to Alexa Fluor 680 (Jacskon ImmunoResearch). Single-stack images were taken at 60x and analyzed in FIJI (ImageJ). pRPA S33, PML, and eGFP-DAXX foci within nuclei (masked according to DAPI staining) were detected with FIJI’s *Find Maxima* function using appropriate thresholds that remained the same for all conditions. The number of pRPA, eGFP-DAXX, and PML maxima were counted for each cell in all conditions. Each PML maximum was then enlarged to a circle with a diameter of 0.2μm to set a boundary in which pRPA S33 or eGFP-DAXX maxima needed to reside in order to be considered “positive.” The raw percentage of pRPA S33 or eGFP-DAXX-positive PML bodies were calculated by dividing the number of maxima within PML bounds by the total number of maxima in the nucleus. The raw percentages for all cells measured in each condition were averaged in each biological replicate.

#### Quantitative analysis of eGFP-DAXX fold enrichment at PML nuclear bodies

Asynchronous U2OS cells were transiently transfected with eGFP-DAXX plasmid as described above and cultured on coverslips. The next day, cells were incubated with ATR inhibitors for 1 hour, after which coverslips were prepared for immunofluorescence microscopy as described above. Cells were immuno-stained with Mouse Monoclonal Anti-PML primary antibody (Santa Cruz Biotechnology) and anti-Mouse secondary antibody conjugated to Alexa Fluor 680 (Jackson ImmunoResearch). Single-stack images were taken at 60x and analyzed in FIJI (ImageJ). The three brightest PML foci were identified using the *Find Maxima* function. A circle with a diameter of 0.2μm was drawn for each PML maxima, and mean eGFP-DAXX intensity was measured within each of these 3 regions. Nuclei were masked according to DAPI staining, and total nuclear eGFP-DAXX was measured for each cell. To calculate fold enrichment at PML nuclear bodies, the mean eGFP-DAXX intensity at the 3 PML regions were averaged and divided by the total nuclear eGFP-DAXX intensity for each cell.

#### Quantitative analysis of eGFP-DAXX localization

Asynchronous HeLa DAXX KO cells were transiently transfected with eGFP-DAXX plasmid as described above and cultured on coverslips. The next day, cells were incubated with 10µM VE-821 for 1 hour, after which coverslips were prepared for immunofluorescence microscopy as described above. Cells were immuno-stained with Mouse Monoclonal Anti-PML primary antibody (Santa Cruz Biotechnology), Human anti-centromere antisera (ACA, Antibodies Inc.), Goat anti-Rabbit secondary antibody conjugated to Alexa Fluor 594 (Jackson ImmunoResearch), and Donkey anti-Human secondary antibody conjugated to Alexa Fluor 680 (Jackson ImmunoResearch). Z-stacks were taken at 60x for each cell with a range of 4µm at 0.3 µm steps. Images were blinded by a colleague and analyzed in FIJI (ImageJ). For each z-step, eGFP-DAXX foci were identified and assessed for co-localization with PML or ACA by eye. eGFP-DAXX foci co-localizing with both PML and ACA foci were counted as only co-localizing to PML nuclear bodies. The number of eGFP-DAXX foci in each cell was counted, and the percentage of eGFP-DAXX foci at PML or centromere foci were calculated.

#### ChIP-Seq analysis of pATR T1989

Given the challenges associated with mapping short reads to repetitive elements and the potential bias of peak calling to identify DNA damage loci rather than basal levels of protein recruitment, we re-analyzed pATR T1989 ChIP-Seq data from Li et al., 2018 along with its input and IgG controls (SRR6428538, SRR6428539, SRR6428540). We identified the relative enrichment of pATR T1989 at centromeric regions by counting the instance of the sequence motif associated with centromere protein B (CENP-B box; ATTCGTTGGAAACGGGA or reverse complement TCCCGTTTCCAACGAAT) using ($grep -c “ATTCGTTGGAAACGGGA” file.fasta) and ($grep -c “TCCCGTTTCCAACGAAT” file.fasta). The number of reads containing the CENP-B box was then normalized to the total number of sequencing reads to determine relative enrichment.

### QUANTIFICATION AND STATISTICAL ANALYSIS

Immunofluorescence intensities, and live-cell imaging cell counts were measured using Fiji software and quantified using Graphpad PRSM software. Gels were analyzed using GelDoc Go System (BioRad) and Fiji software. Two-tailed student’s t test for replicate averages were performed for all statistical analysis as shown in figure legends.

## SUPPLEMENTAL FIGURE LEGENDS

**Figure S1:**

**(A-B)** *Total CENP-A levels are unaffected by ATR inhibition*. (A) Representative western blot of asynch. U2OS in cells that were untreated (CONT) or treated for 1 hour with 10μM VE-821 or 10μM AZ20 (ATRi). (B) Quantification of mean nuclear CENP-A intensity in asynch. U2OS cells that were treated as in (A). (>100 cells per condition; n=3 biological replicates). Error bars represent mean ±SD.

**(C-D)** *ATR inhibition induces the targeted loss of CENP-A nucleosomes from interphase centromeres*. Quantitative PCR of CENP-A (C) or H2A (D) chromatin immunoprecipitation in asynch. U2OS cells treated as in (A). Centromeres were amplified using primers for the core α-satellite sequences of chromosome 4 (αSat-4) or chromosomes 1, 5, and 19 (αSat-1). Error bars represent mean ±SEM.

**(E)** *ATR activity is rapidly depleted following treatment with VE-821*. Representative western blot of asynch. U2OS cells treated for 18 hours with 0.3μM Aphidicolin to increase ATR activity, then cells were treated with 10μM VE-821 (ATRi) for 30 minutes. Cells were collected for western blotting at 10, 20, and 30 mins after ATRi for analysis of pChk1 (S345) levels.

*p ≤ 0.05, **p ≤ 0.01, two-tailed t-test of replicate averages.

**Figure S2:**

**(A)** *Asynchronous cells are sorted into their distinct cell cycle phases according to EdU and DAPI staining*. Representative cell cycle plots of asynch. U2OS cells that were untreated (CONT) or treated for 1 hour with with 10μM VE-821 or 10μM AZ20.

**(B-C)** *Acute ATR inhibition does not increase DNA damage*. (B) Representative western blot of asynch. U2OS cells that were untreated (CONT) or treated for 1 hour with 10μM VE-821, 10μM AZ20, or 0.5 mM Doxorubicin (DOXO). (C) Quantification of mean nuclear γH2AX intensity in asynch. U2OS cells that were treated as in (A). (>50 cells per condition; n = 3 biological replicates). Outliers were removed according to the ROUT (Q=1%) method. Error bars represent mean ±SD.

**(D-E)** *CENP-A loss is not induced by acute Chk1 or ATM inhibition*. Quantification of maximum nuclear CENP-A intensity in asynch. U2OS cells that were untreated (CONT) or treated for 1 hour with Chk1 inhibitor (2 µM MK8776) or ATM inhibitor (10 µM KU 55933). (>50 cells per condition; n = 3 biological replicates). Outliers were removed according to the ROUT (Q=1%) method. Error bars represent mean ±SD.

*p ≤ 0.05, **p ≤ 0.01, two-tailed t-test of replicate averages.

**Figure S3:**

**(A-B)** *ATR activity is not enriched at interphase centromeres*. (A) ChIP-Seq Analysis of pATR T1989 data from Li et al., 2018 for enrichment at sequencing reads containing the CENP-B box. (B) Quantification of the percentage of pRPA S33 foci in untreated asynch. U2OS cells that localize to PML nuclear bodies or centromeres marked by anti-centromere antibody (ACA). (>280 cells per condition; n=3 biological replicates). Error bars represent mean ±SEM.

**(C)** *ATR inhibition reduces the proportion of DAXX-positive PML nuclear bodies*. Asynch. U2OS cells were untreated (CONT) or treated for 1 hour with 10μM VE-821 or 10μM AZ20 (ATRi). (>50 cells per condition; n = 3 biological replicates). Error bars represent mean ±SEM.

**(D)** *HeLa DAXX KO cells are sensitive to ATR inhibitors*. Representative western blot of asynch. HeLa WT (parental) or DAXX KO cells that were untreated (CONT) or treated for 1 hour with 10μM VE-821 or 10μM AZ20.

**(E-F)** *ATR inhibition increases DAXX and H3*.*3 occupancy at interphase centromeres*. Quantitative PCR of DAXX or H3.3 chromatin immunoprecipitation in asynch. U2OS cells treated as in (C). Centromeres were amplified using primers for the core α-satellite sequences of chromosomes 1, 5 and 19 (αSat-1) or chromosome 4 (αSat-4). Error bars represent mean ±SEM.

**(G)** *DAXX residues matching ATR’s consensus motif are phosphorylated near the C-terminal SIM*. Domain map of human DAXX protein with SIMs indicated. S/T-Q phosphorylation identified using PhosphoSitePlus are mapped accordingly. Residues of interest are highlighted in blue.

**(H)** *Phospho-null mutation of eGFP-DAXX decreases maximum nuclear CENP-A intensity and de-sensitizes cells to ATR inhibition*. Asynch. HeLa DAXX KO cells were untransfected (-) or transiently transfected with eGFP-DAXX^WT^ or eGFP-DAXX^AA^ (S707A;S712A). Cells were untreated (-) or treated for 1 hour with 10μM VE-821 (+ATRi) and stained for CENP-A (red) and PML (magenta). Scale bar = 5 μm.

*p ≤ 0.05, **p ≤ 0.01, two-tailed t-test of replicate averages.

**Figure S4:**

**(A)** *G1 U2OS cells in the following cell cycle are isolated after acute ATR inhibition without DNA damage induction*. Asynchronous cells were untreated (CONT) or treated for 1 hour with 10μM VE-821, after which VE-821 was washed out. Cells were then incubated with Nocodazole (Noc; 50 ng/mL) for 16 hours, after which Noc was washed out and G1 cells were collected for western blotting. The absence of Cyclin A indicates that G1 cells have not progressed into S-phase.

**(B-C)** *ATR activity is restored after inhibitor wash-out in HeLa WT and DAXX KO cells*. Representative western blot of asynch. HeLa WT (parental) and HeLa DAXX KO cells that were untreated (CONT) or treated for 1 hour with 10μM VE-821 (ATRi), after which VE-821 was washed out. Cells were collected for western blotting 6 hours after wash-out.

**(D-E)** *ATR inhibition increases DAXX localization to peri-centromeres and telomeres*. Quantitative PCR of DAXX chromatin immunoprecipitation in asynch. U2OS cells that were untreated (CONT) or treated for 1 hour with 10μM VE-821 or 10μM AZ20 (ATRi). (D) Centromeres were amplified using primers for pericentromeric Sat2 DNA sequences (Sat2). (E) Telomeres were amplified using primers for the sub-telomeric regions of chromosomes 1 and 22 (Sub.Tel). Error bars represent mean ±SEM.

*p ≤ 0.05, **p ≤ 0.01, two-tailed t-test of replicate averages.

## Notes

### Competing Interest Statement

The authors have declared no competing interest.

