## Supplemental Figures and Figure Legends for "ATR protects centromere identity by promoting DAXX association with PML nuclear bodies"

*p ≤ 0.05, **p≤ 0.01, two-tailed t-test of replicate averages.

**Supplemental Figure 3:**

**(A-B)** *ATR activity is not enriched at interphase centromeres.* (A) ChIP-Seq Analysis of pATR T1989 data from Li et al., 2018 for enrichment at sequencing reads containing the CENP-B box.

(B) Quantification of the percentage of pRPA S33 foci in untreated asynch. U2OS cells that localize to PML nuclear bodies or centromeres marked by anti-centromere antibody (ACA). (>280 cells per condition; n=3 biological replicates). Error bars represent mean $\pm$ SEM.

**(C)** *ATR inhibition reduces the proportion of DAXX-positive PML nuclear bodies.* Asynch. U2OS cells were untreated (CONT) or treated for 1 hour with 10μM VE-821 or 10μM AZ20 (ATRi). (>50 cells per condition; n = 3 biological replicates). Error bars represent mean $\pm$ SEM.

**(D)** *HeLa DAXX KO cells are sensitive to ATR inhibitors.* Representative western blot of asynch. HeLa WT (parental) or DAXX KO cells that were untreated (CONT) or treated for 1 hour with 10μM VE-821 or 10μM AZ20.

**(E-F)** *ATR inhibition increases DAXX and H3.3 occupancy at interphase centromeres.* Quantitative PCR of DAXX or H3.3 chromatin immunoprecipitation in asynch. U2OS cells treated as in (C). Centromeres were amplified using primers for the core α-satellite sequences of chromosomes 1, 5 and 19 (αSat-1) or chromosome 4 (αSat-4). Error bars represent mean $\pm$ SEM.

**(G)** *DAXX residues matching ATR's consensus motif are phosphorylated near the C-terminal SIM.* Domain map of human DAXX protein with SIMs indicated. S/T-Q phosphorylation identified using PhosphoSitePlus are mapped accordingly. Residues of interest are highlighted in blue.

**(H)** *Phospho-null mutation of eGFP-DAXX decreases maximum nuclear CENP-A intensity and de-sensitizes cells to ATR inhibition*. Asynch. HeLa DAXX KO cells were untransfected (-) or transiently transfected with eGFP-DAXX$^{WT}$ or eGFP-DAXX$^{AA}$ (S707A;S712A). Cells were untreated (-) or treated for 1 hour with 10μM VE-821 (+ATRi) and stained for CENP-A (red) and PML (magenta). Scale bar = 5 μm.

*p $\leq$ 0.05, **p$\leq$ 0.01, two-tailed t-test of replicate averages.

**Supplemental Figure 4:**

**(A)** *G1 U2OS cells in the following cell cycle are isolated after acute ATR inhibition without DNA damage induction.* Asynchronous cells were untreated (CONT) or treated for 1 hour with 10μM VE-821, after which VE-821 was washed out. Cells were then incubated with Nocodazole (Noc; 50 ng/mL) for 16 hours, after which Noc was washed out and G1 cells were collected for western blotting. The absence of Cyclin A indicates that G1 cells have not progressed into S-phase.

**(B-C)** *ATR activity is restored after inhibitor wash-out in HeLa WT and DAXX KO cells.* Representative western blot of asynch. HeLa WT (parental) and HeLa DAXX KO cells that were untreated (CONT) or treated for 1 hour with 10μM VE-821 (ATRi), after which VE-821 was washed out. Cells were collected for western blotting 6 hours after wash-out.

**(D-E)** *ATR inhibition increases DAXX localization to peri-centromeres and telomeres.* Quantitative PCR of DAXX chromatin immunoprecipitation in asynch. U2OS cells that were untreated (CONT) or treated for 1 hour with 10μM VE-821 or 10μM AZ20 (ATRi). (D) Centromeres were amplified using primers for pericentromeric Sat2 DNA sequences (Sat2). (E) Telomeres were amplified using primers for the sub-telomeric regions of chromosomes 1 and 22 (Sub.Tel). Error bars represent mean $\pm$ SEM.

*p $\leq$ 0.05, **p$\leq$ 0.01, two-tailed t-test of replicate averages.

**Figure S1**

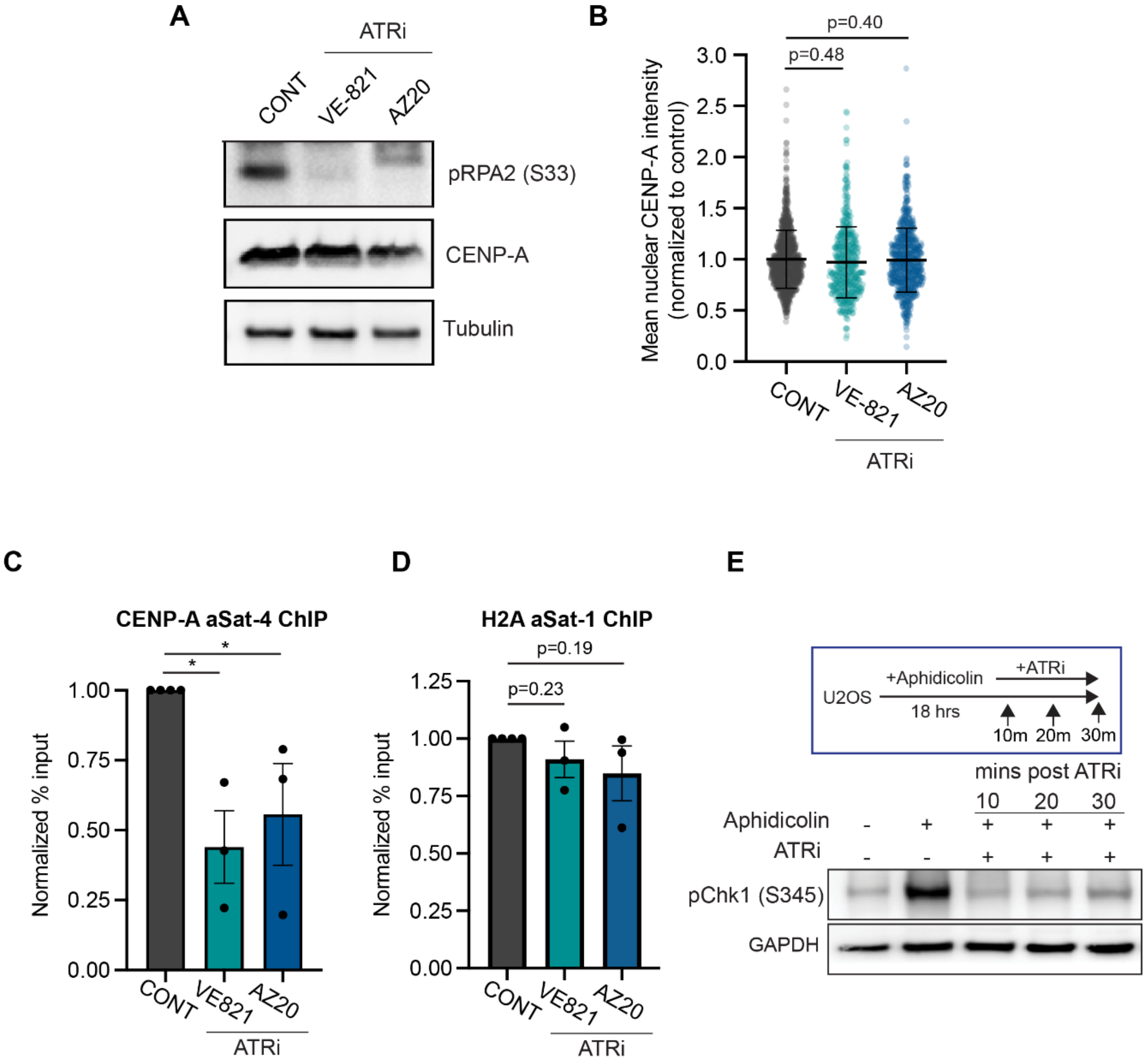
A
ATRi
CONT
VE-821
AZ20
pRPA2 (S33)
CENP-A
Tubulin
B
p=0.40
p=0.48
Mean nuclear CENP-A intensity (normalized to control)
3.0
2.5
2.0
1.5
1.0
0.5
0.0
CONT
VE-821
AZ20
ATRi
C
CENP-A aSat-4 ChIP
*
*
Normalized % input
1.00
0.75
0.50
0.25
0.00
CONT
VE821
AZ20
ATRi
D
H2A aSat-1 ChIP
p=0.19
p=0.23
Normalized % input
1.25
1.00
0.75
0.50
0.25
0.00
CONT
VE821
AZ20
ATRi
E
+ATRi
+Aphidicolin
U2OS
18 hrs
10m 20m 30m
mins post ATRi
10 20 30
Aphidicolin - + + + +
ATRi - - + + +
pChk1 (S345)
GAPDH

**Figure S2**

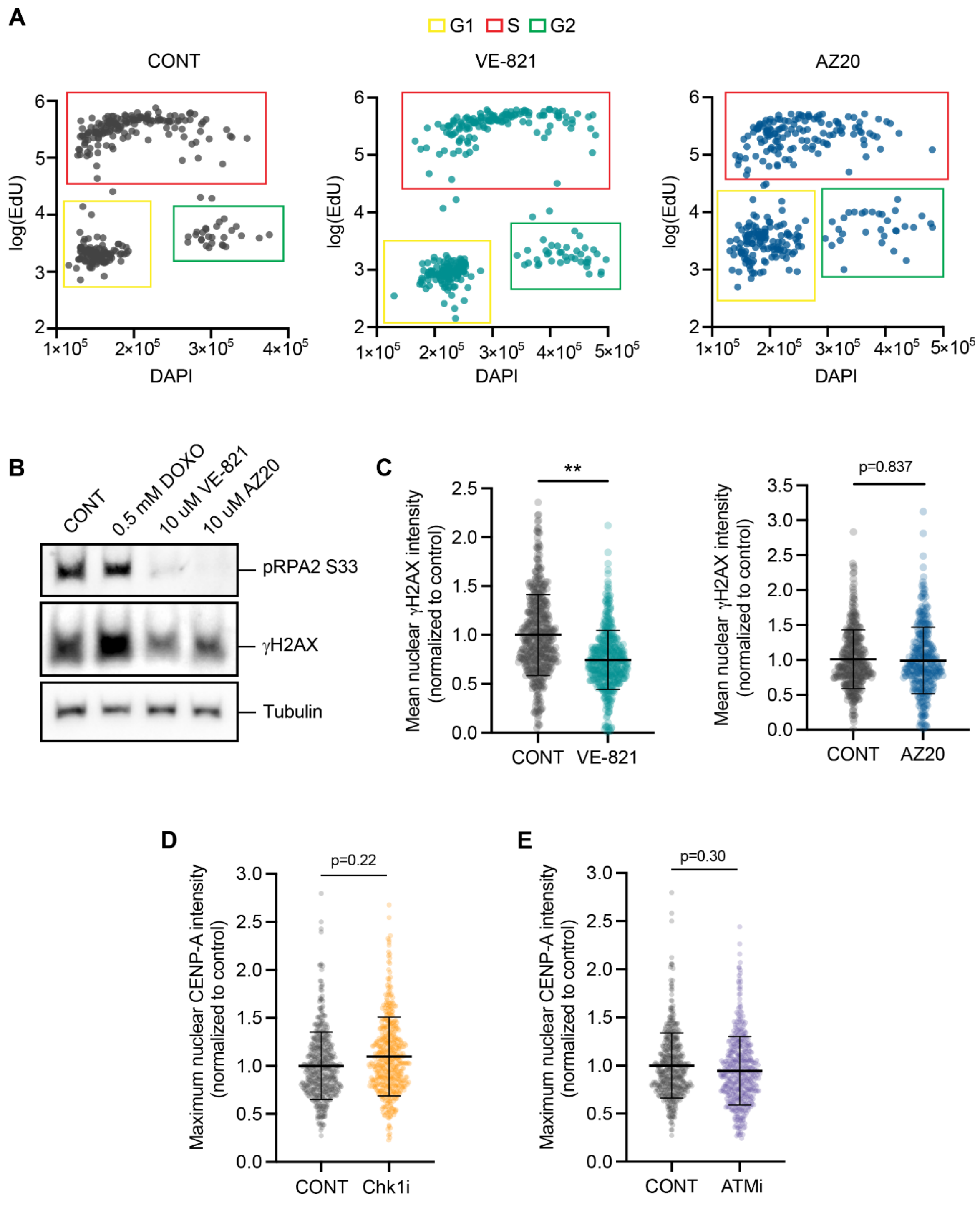

A
G1
S
G2
CONT
VE-821
AZ20
log(EdU)
DAPI
B
CONT
0.5 mM DOXO
10 uM VE-821
10 uM AZ20
pRPA2 S33
γH2AX
Tubulin
C
Mean nuclear γH2AX intensity (normalized to control)
**
p=0.837
CONT
VE-821
AZ20
D
Maximum nuclear CENP-A intensity (normalized to control)
p=0.22
CONT
Chk1i
E
p=0.30
CONT
ATMi

### Figure S3

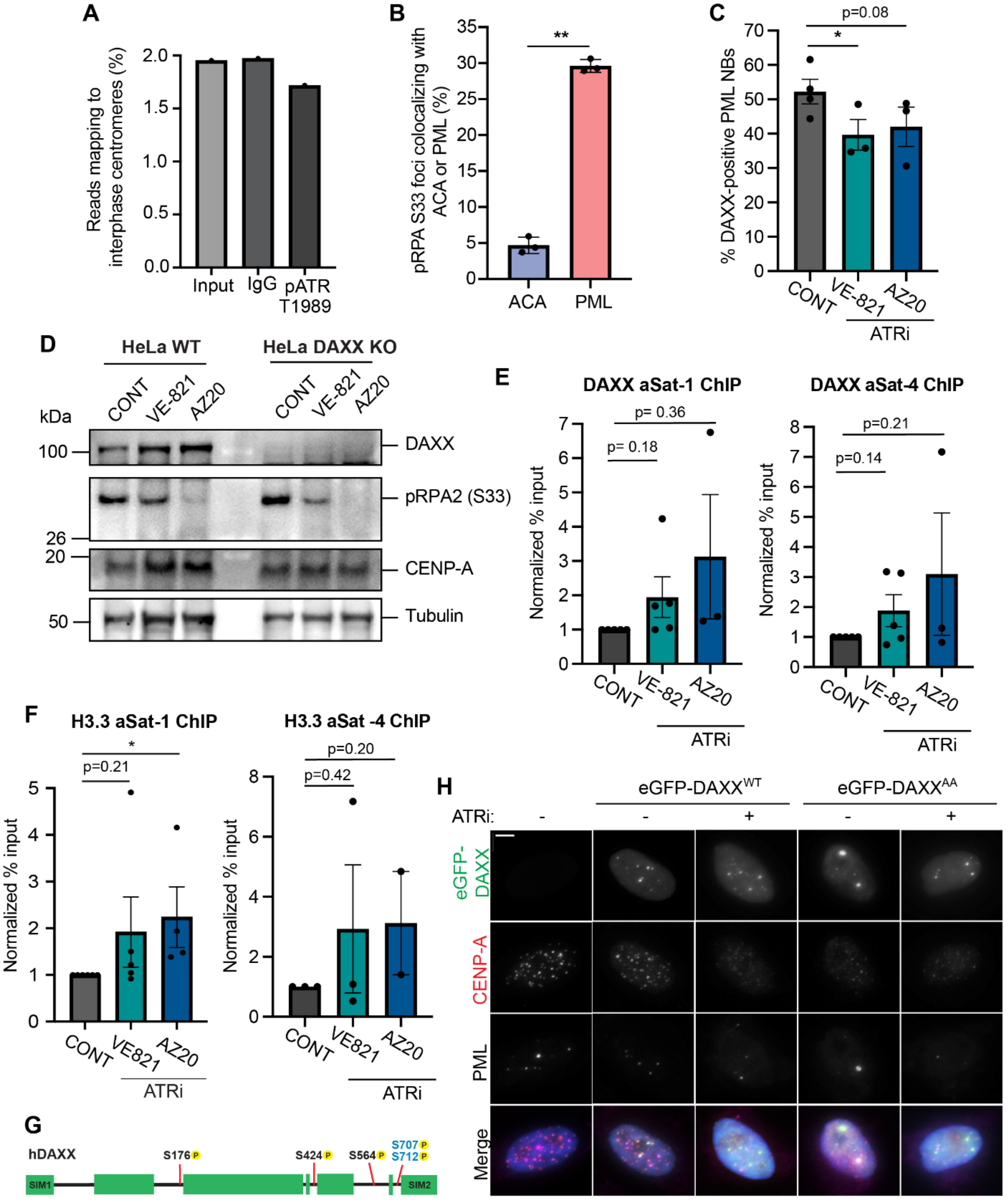

A
Reads mapping to interphase centromeres (%)
2.0
1.5
1.0
0.5
0.0
Input
IgG
pATR T1989
B
pRPA S33 foci colocalizing with ACA or PML (%)
35
30
25
20
15
10
5
0
**
ACA
PML
C
% DAXX-positive PML NBs
70
60
50
40
30
20
10
0
p=0.08
*
CONT
VE-821
AZ20
ATRi
D
HeLa WT
HeLa DAXX KO
CONT
VE-821
AZ20
kDa
100
26
20
50
DAXX
pRPA2 (S33)
CENP-A
Tubulin
E
DAXX aSat-1 ChIP
DAXX aSat-4 ChIP
Normalized % input
p= 0.36
p= 0.18
p=0.21
p=0.14
CONT
VE-821
AZ20
ATRi
F
H3.3 aSat-1 ChIP
H3.3 aSat -4 ChIP
Normalized % input
*
p=0.21
p=0.20
p=0.42
CONT
VE821
AZ20
ATRi
G
hDAXX
S176 P
S424 P
S564 P
S707 P
S712 P
SIM1
SIM2
H
eGFP-DAXXWT
eGFP-DAXXAA
ATRi:
-
+
eGFP-DAXX
CENP-A
PML
Merge

### Figure S4

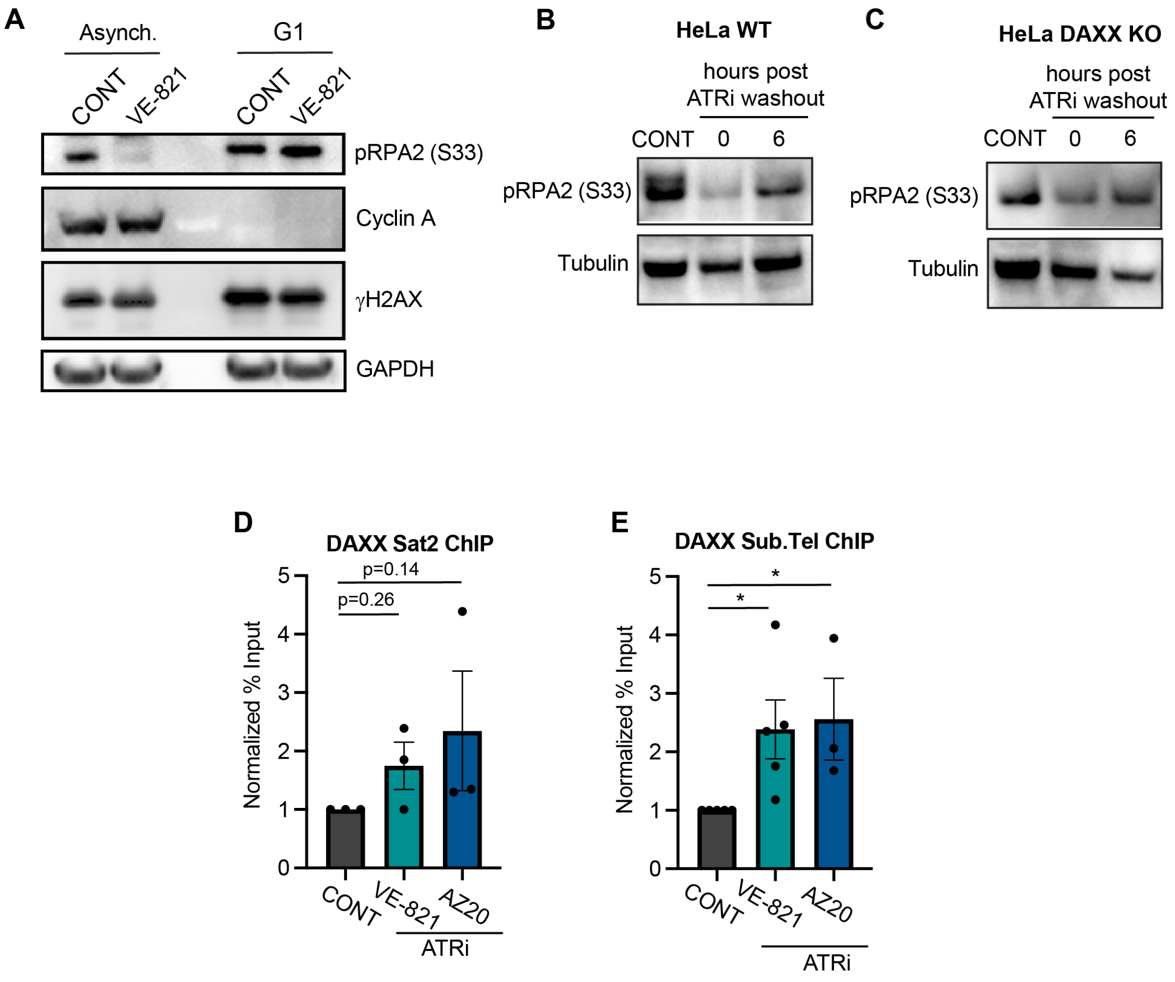
A
Asynch.
G1
CONT
VE-821
CONT
VE-821
pRPA2 (S33)
Cyclin A
γH2AX
GAPDH
B
HeLa WT
hours post ATRi washout
CONT
0
6
pRPA2 (S33)
Tubulin
C
HeLa DAXX KO
hours post ATRi washout
CONT
0
6
pRPA2 (S33)
Tubulin
D
DAXX Sat2 ChIP
p=0.14
p=0.26
Normalized % Input
0
1
2
3
4
5
CONT
VE-821
AZ20
ATRi
E
DAXX Sub.Tel ChIP
*
*
Normalized % Input
0
1
2
3
4
5
CONT
VE-821
AZ20
ATRi
